# Single-Cell Atlas of AML Reveals Age-Related Gene Regulatory Networks in t(8;21) AML

**DOI:** 10.1101/2024.10.29.620871

**Authors:** Jessica Whittle, Stefan Meyer, Georges Lacaud, Syed Murtuza Baker, Mudassar Iqbal

## Abstract

**Background:** Acute myeloid leukemia (AML) is characterized by cellular and genetic heterogeneity, which correlates with clinical course. Although single-cell RNA sequencing (scRNA-seq) reflects this diversity to some extent, the low sample numbers in individual studies limit the analytic potential when comparing specific patient groups.

**Results:** We performed large scale integration of published scRNA-seq datasets to create a unique single-cell transcriptomic atlas for AML (AML scAtlas), totaling 748,679 cells, from 159 AML patients and 44 healthy donors from 20 different studies. This is the largest single-cell data resource for AML to our knowledge, publicly available at https://cellxgene.bmh.manchester.ac.uk/AML/. This AML scAtlas allowed investigations into 20 patients with t(8;21) AML, where we explored the clinical importance of age, given the *in-utero* origin of pediatric disease. We uncovered age-associated gene regulatory network (GRN) signatures, which we validated using bulk RNA sequencing data to delineate distinct groups with divergent biological characteristics. Furthermore, using an additional multiomic dataset (scRNA-seq and scATAC-seq), we validated our initial findings and created a de-noised enhancer-driven GRN reflecting the previously defined age-related signatures.

**Conclusions:** Applying integrated data analysis of the AML scAtlas, we reveal age-dependent gene regulation in t(8;21) AML, potentially reflecting immature/fetal HSC origin in prenatal origin disease vs postnatal origin. Our analysis revealed that BCLAF1, which is particularly enriched in pediatric AML with t(8;21) of inferred *in-utero* origin, is a promising prognostic indicator. The AML scAtlas provides a powerful resource to investigate molecular mechanisms underlying different AML subtypes.

## Introduction

Acute myeloid leukemia (AML) is an aggressive blood cancer driven by non-random genomic rearrangements in hematopoietic stem/progenitor cells (HSPCs). Recurrent AML-associated genomic aberrations, which often involve transcriptional or epigenetic regulators, give rise to distinct patterns of gene expression strongly associated with clinical course and chemotherapy response ^[1, 2]^. Single-cell RNA sequencing (scRNA-seq) studies have demonstrated that HSPCs acquire lineage priming at an early stage when still phenotypically immature and disperse down an erythromyeloid or lymphomyeloid differentiation trajectory ^[3]^. In the context of AML, diverse clonal hierarchies include the co-existence of normal hematopoietic clones. Leukemic clones can partially recapitulate myeloid differentiation and have been shown to display functional differences even when defined by the same genotype ^[4-7]^. Indeed, analysis of AML using scRNA-seq has revealed key clonal hierarchies, defining subtype-associated cell types, and dynamic changes following therapy, and have been critical in characterizing leukemic stem cells (LSCs), which propagate the disease and drive relapse ^[5-8]^.

Most AML scRNA-seq studies are limited by small sample numbers and include a mixture of different AML subtypes which may not be directly comparable to one another. Therefore, it is difficult to make biological conclusions with sufficient robustness to be clinically translatable in these individual datasets. To overcome this, we performed large-scale integration of public scRNA-seq datasets to create a single-cell transcriptomic atlas for AML (AML scAtlas). Due to the range of data sources spanning time, locations, and experimental designs, complex batch effects often arise between scRNA-seq datasets which requires a tailored data integration approach ^[9, 10]^. Thus, we benchmarked some widely used batch correction tools ^[11-13]^ for our specific data use case.

Given the broad representation of age groups in AML scAtlas, we sought to investigate a developmental aspect of AML biology. Pediatric AMLs have substantially better clinical outcomes compared to adult AMLs^[14-16]^. The molecular landscape of AML differs between children and adults^[2, 14-16]^; this may, in part, reflect differences in the developmental origins of the disease. Chromosomal changes in pediatric leukemia are acquired *in-utero*, as evidenced by leukemia-specific genomic aberrations detected in the Gunthrie spots of children who later developed leukemia, sometimes several years after birth ^[17]^. Adult leukemia, in contrast, is thought to develop later in life through acquisition of pre-leukaemic changes and clonal evolution of adult HSPCs ^[18, 19]^. The impact of these developmental stages on leukemia biology remains incompletely understood, and no current methods exist to quantify and characterize differences in the origin of the disease. However, as childhood AML with presumed *in-utero* origin has a better outcome, for teenagers and young adults, determination of the pre- or postnatal origin might be important for better treatment stratification and prognostication.

AML with t(8;21) (AML-ETO/RUNX1-RUNX1T1) is one of the most frequent AML subtypes in young people, although it affects all ages ^[2]^. The prenatal origins of the t(8;21) rearrangement, has been confirmed even in older children presenting with AML ^[17]^. The prognosis of AML with t(8;21) is better in children than in teenagers and even more so than in young adults ^[20]^. This outcome difference is not fully explainable by co-morbidities and may instead be related to the developmental origins of the disease. In the intermediate teenage group, t(8;21) AML may comprise both late childhood and early adult disease entities, a distinction that could have prognostic implications and could help to explain disease biology and clinical course.

We leveraged our AML scAtlas resource to characterize age and developmental stage specific signatures in t(8;21) AML by applying single-cell gene regulatory network (GRN) inference^[21, 22]^, as a means of revealing cell state heterogeneity across age groups. We then validated and refined our findings in a larger cohort using bulk RNA sequencing (RNA-seq) data from the TARGET^[2]^ and BeatAML^[23]^ studies, defining age-associated GRN signatures and key regulators of t(8;21) AML, that may reflect the developmental origins of the leukemia.

Profiling both gene expression and chromatin accessibility together can decipher the enhancer-driven GRN (eGRN) and enriched transcriptional regulators. Significant heterogeneity across different patients and time points ^[24]^ was recently described by analyzing combined scRNA-seq and single-cell Assay for Transposase Accessible Chromatin sequencing (scATAC-seq). We used the t(8;21) AML data from this study to validate our initial findings, by applying cutting edge GRN inference methodology ^[25]^. This encompasses both modalities to provide a denoised eGRN which we could correlate with our age-associated signatures.

## Results

### Large Scale Data Integration to Construct a Single-Cell Transcriptomic Atlas of AML (AML scAtlas)

To create the AML scAtlas, we integrated published scRNA-seq data of primary AML bone marrow samples, from 16 suitable high-quality studies (**see Materials and Methods**), comprising 159 AML samples (**Figure 1A; Supplementary Table 1**). Where on-treatment time points were available, we selected only diagnostic samples to establish a reference atlas of primary AML at diagnosis. If studies had healthy donor bone marrow samples, these were included, alongside data from healthy bone marrow samples from four additional scRNA-seq studies (**Supplementary Table 1**) to enable comparisons between malignant and healthy bone marrow populations. After cell filtering and quality control, the AML scAtlas contains data from 748,679 high quality cells derived from a total of 20 different scRNA-seq studies ^[4-6, 8, 26-40]^ (**Supplementary Figure 1A**). Each sample was assigned to an AML clinical subtype, based on the recent European Leukemia Net (ELN) clinical guidelines ^[41]^, and classified into the corresponding prognostic risk group. This resource captures a broad range of molecular subtypes of AML and spans different age groups, including both pediatric and adult AML cases (**Figure 1B;1C**). Overall, this is the largest dataset to date for exploring AML biology at single-cell resolution.

**Figure 1.**
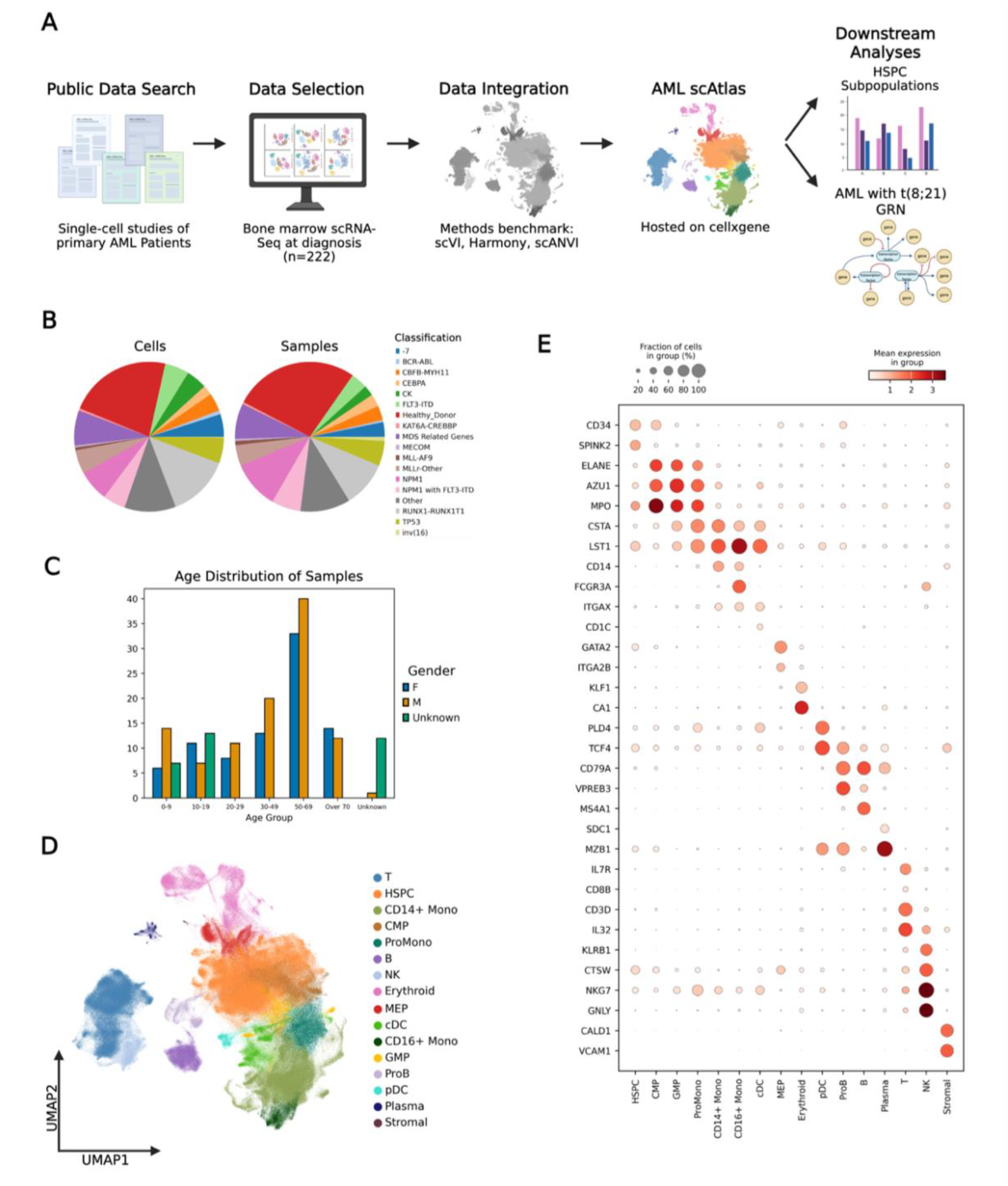
Large Scale Data Integration Creates a Single-Cell Atlas of AML. (**A**) Overview of the analysis steps in creating AML scAtlas. (**B**) Proportion of cells (left panel) and samples (right panel) belonging to each AML subtype as defined by the ELN clinical guideline. (**C**) Age group and gender distribution of AML scAtlas cohort samples. (**D**) scVI harmonized UMAP colored by annotated cell types. (**E**) The expression of key hematopoietic marker genes across annotated cell types shown on a dotplot. Color scale shows mean gene expression, dot size represents the fraction of cells expressing the given gene.

In the initial analysis of the combined dataset batch effects were noted, with study-specific clustering, which was quantified using several benchmarking metrics (**Supplementary Table 2; Supplementary Figure 1A;1B**). Even within samples of the same study, sample-wise clustering was noted (**Supplementary Figure 1C**). To address this, we benchmarked several widely used batch correction methods (**Supplementary Table 2**; **Supplementary Figure 2A;2C**), identifying scVI as the best method for this dataset (**Supplementary Table 2**). We therefore employed scVI to correct for batch effects, before clustering and cell type annotation in the AML scAtlas, by using the consensus of multiple annotation tool results (**Supplementary Table 3**), verified using cluster-wise marker gene expression (**Figure 1D;1E**).

**Figure 2.**
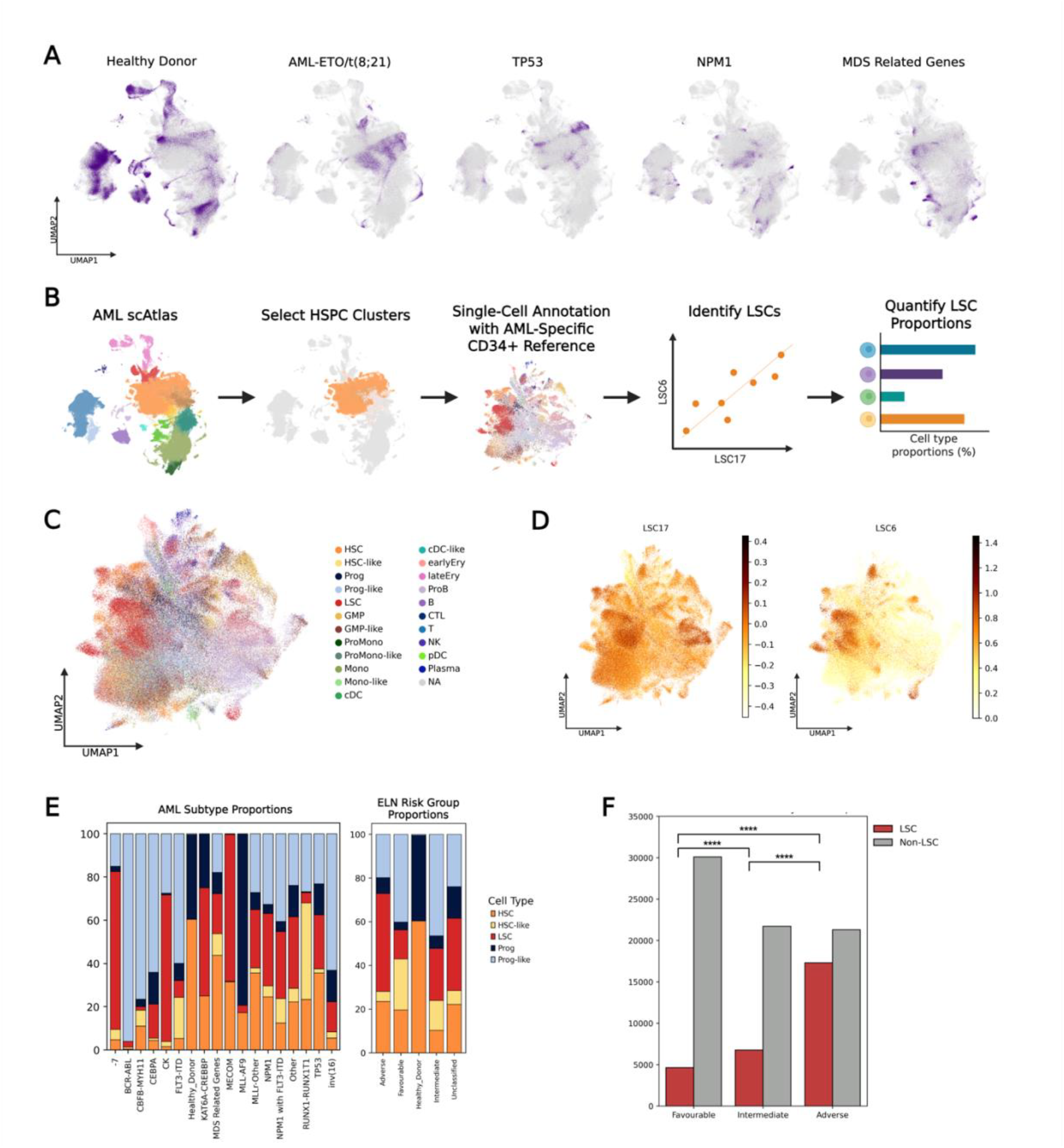
Characterizing Cell Type Distributions in AML Subtypes. (**A**) UMAP highlighting the distribution of cells from different AML subtypes in AML scAtlas. (**B**) Schematic showing the workflow used to identify leukemic stem cells (LSCs) from the AML scAtlas hematopoietic stem and progenitor cell (HSPC) clusters. (**C**) Using the AML scAtlas HSPC clusters only, UMAP was regenerated and annotated with an AML-specific reference of leukemia stem and progenitor cells (LSPCs). (**D**) UMAPs showing the leukemic stem cell scores of each cell, for the LSC17 (left) and LSC6 (right). (**E**) Proportions of HSPC/LSPC populations in different AML subtypes (left) and AML risk groups (right), as defined by ELN clinical guidelines. (**F**) Comparison of LSC abundance in favourable and adverse ELN risk groups. Chi-Square test statistic: 8658.98, degrees of freedom: 1, P-value: 0.0.

Cell type proportions analyses across the clinically relevant subtypes in the dataset show that the AML subtypes were significantly biased towards myeloid cell types (CMP, MEP, GMP, ProMono, CD14+ Mono, CD16+ Mono, cDC, Erythroid) with each subtype exhibiting a clear predominant cell type consistent with AML clonal expansion (**Figure 2A; Supplementary Figure 3A**). In contrast, healthy donor samples had more balanced lineage proportions, with lymphoid cells (T, B, NK, ProB, pDC, Plasma) well represented (**Figure 2A; Supplementary Figure 3A**). Given the established critical role of HSPCs and LSCs in propagating AML, and their importance as therapeutic targets ^[42]^, we focused on HSPC clusters for further analysis (**Figure 2B**). To identify LSCs, we applied a curated reference profile of leukaemic stem and progenitor cells (LSPCs) ^[7]^ (**Figure 2B;2C**) and correlated this with calculated LSC6 ^[43]^ and LSC17^[44]^ scores for each cell (**Figure 2D**). We then compared the proportions of HSPC/LSPCs across different AML subtypes and risk groups, as defined by the ELN clinical guidelines ^[41]^ (**Figure 2E**). Higher-risk subtypes displayed a higher proportion of LSCs compared to favorable risk disease (**Figure 2E;2F**).

**Figure 3.**
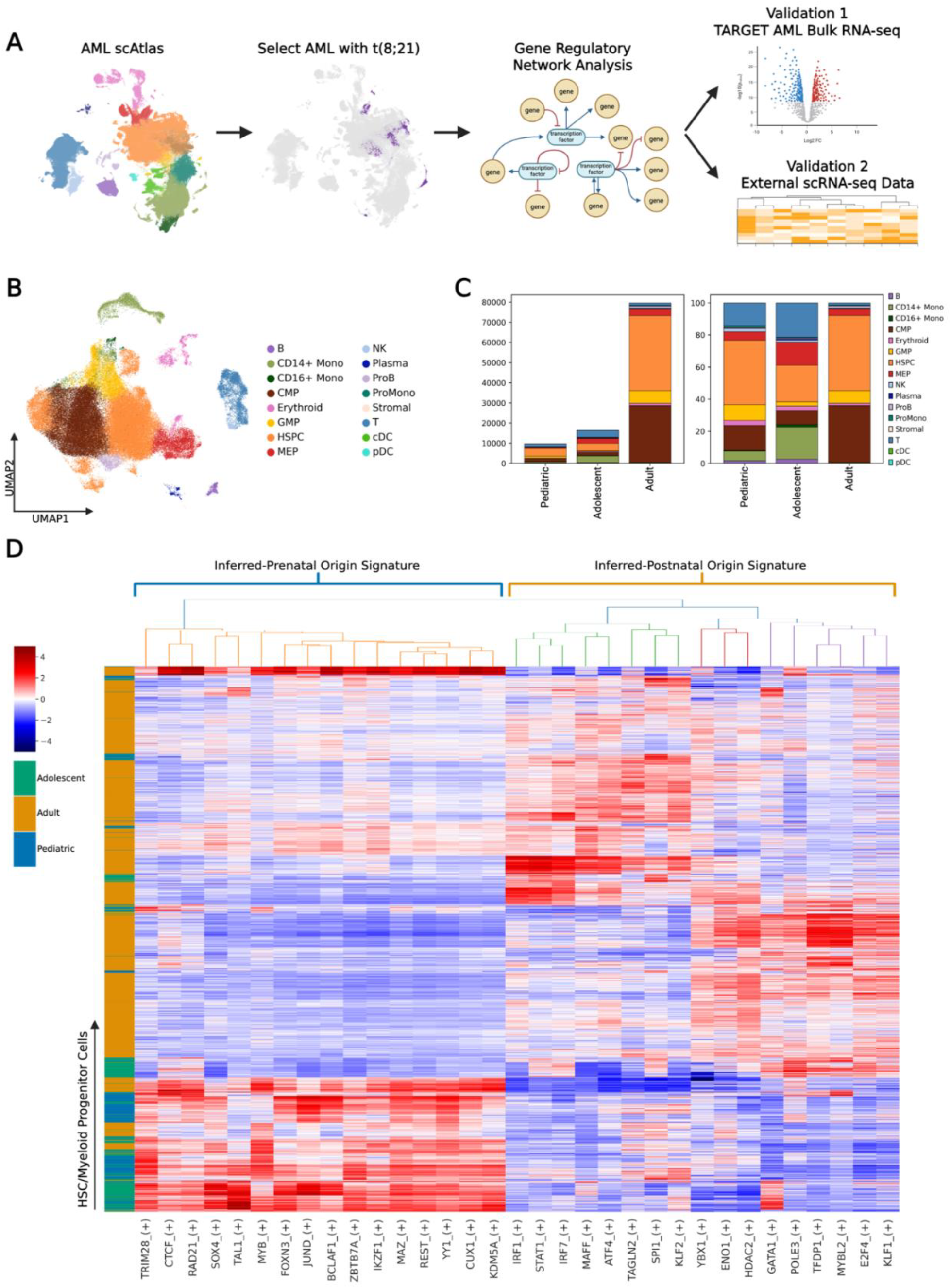
AML scAtlas Reveals Age-Associated Heterogeneity in t(8;21) AML. (**A**) Depiction of the workflow to generate and validate the t(8;21) AML gene regulatory network (GRN) from AML scAtlas. (**B**) Using the AML scAtlas t(8;21) sample cells, UMAP was re-computed and shows the different cell types. **(C)**Bar plots of the absolute cell type numbers (left panel) and the cell type proportions (right panel) stratified by age group. The CD34 enrichment performed on several adult samples is reflected. (**D**) Using HSPCs and CMPs only, the pySCENIC gene regulatory network (GRN) and regulon AUC scores were calculated. Z-score normalized scores underwent hierarchical clustering to create a clustered heatmap and identify age-associated regulons. Regulons were prioritized using their regulon specificity scores (RSS).

### Application of AML scAtlas to Identifying Age-Associated Gene Regulatory Networks in t(8;21) AML

The AML scAtlas enables robust comparison of adult and pediatric AML. We hypothesized that in adolescents and young adults with t(8;21) AML, the potential for either *in-utero* or postnatal HSPC origin disease might affect disease biology and prognosis. Thus, we sought to explore biological differences between pediatric and adult cases of t(8;21) AML, aiming to explain and potentially improve prognostication in adolescents and young adults. We selected samples with t(8;21) AML from the AML scAtlas, resulting in 105,663 cells from 13 adult cases (aged 20-67), 7 adolescent cases (aged 12-17), and 3 pediatric cases (aged 6-8) (**Figure 3A-3C**). Where gender information was not available, this was inferred from ChrY/XIST gene expression (**Supplementary Figure 4A**). Several adult samples underwent CD34 selection in original studies, excluding more differentiated cell types (mature lymphoid populations, monocytes, granulocytes) in these samples. Thus, these cell types were excluded from comparative analysis, focusing only on HSPCs and myeloid progenitors (CMP, GMP, MEP), which were well represented in all studies (**Figure 3C**).

**Figure 4.**
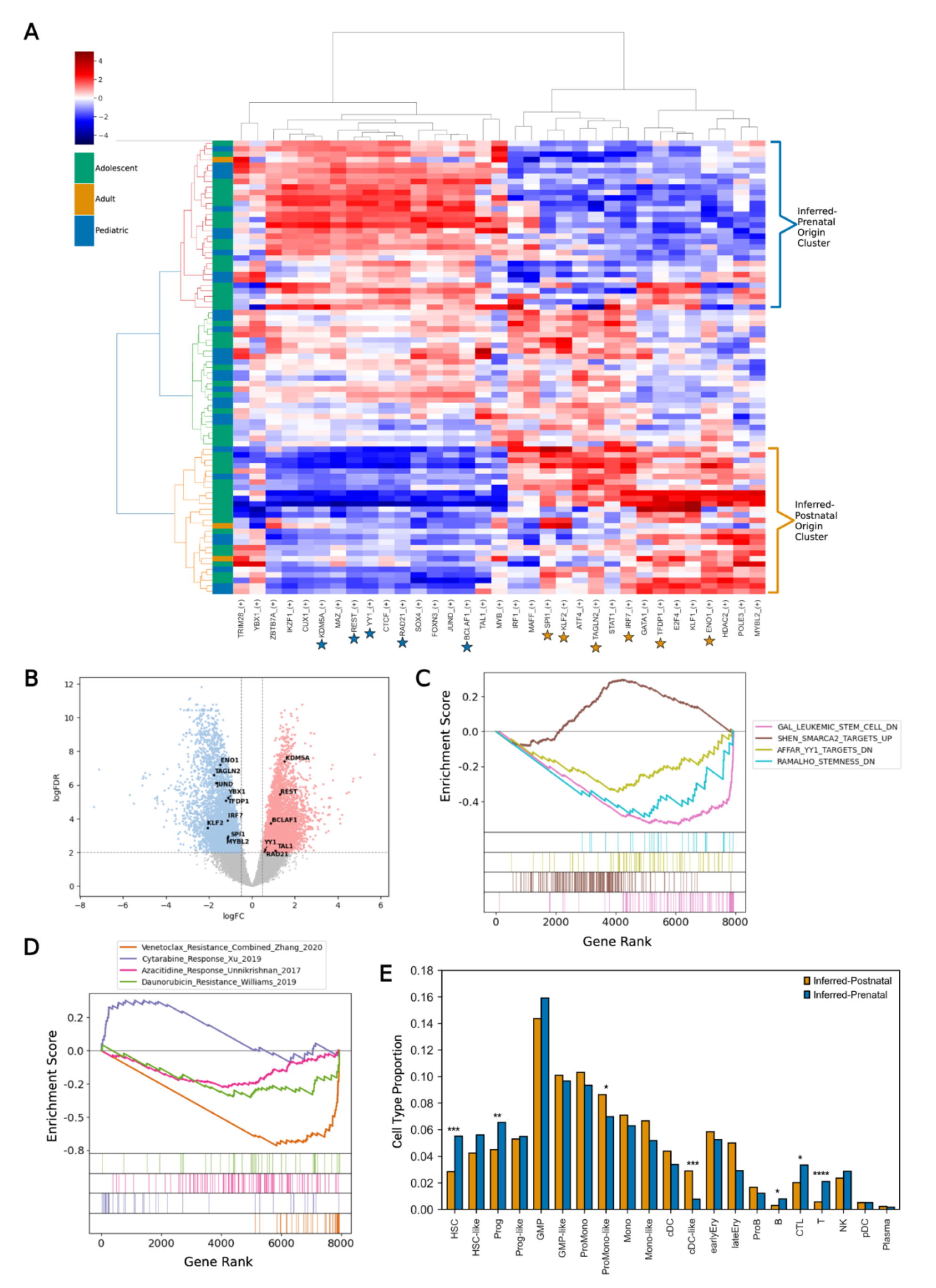
Validation of Age-Associated Regulons in Large Bulk RNA-Seq Cohorts. **(A)** Using previously defined age-associated regulons, pySCENIC AUC scores (Z-score normalized) were clustered to identify samples most enriched for inferred-prenatal and inferred-postnatal origin signatures. (**B**) Volcano plot of differentially expressed genes when comparing the inferred-prenatal origin and inferred-postnatal origin samples. Adjusted P value threshold 0.01; log2 fold change threshold 0.5. Regulon signature associated TFs are indicated. (**C**) Enrichment plot of significant gene sets enriched in the inferred-prenatal origin samples. GSEA was performed on the DEGs using MSigDB databases. FDR q-value threshold <0.05. (**D**) Enrichment plot of drug sensitivity gene sets enriched in the inferred-prenatal samples. GSEA was performed on the DEGs, using drug response signatures from published studies of 4 widely used AML drugs. FDR q-value threshold <0.05. (**E**) The predicted cell type proportions estimated using AutoGeneS deconvolution, of the inferred-prenatal and inferred-postnatal origin samples were compared using T-Tests. Significant P values <0.05 (*), <0.01 (**), <0.001 (***) and <0.0001 (****) are indicated.

We reconstructed the GRN for the t(8;21) subset using the pySCENIC ^[22]^ pipeline, which is a python-based efficient implementation of original SCENIC method ^[21]^. It is a state-of-the-art method for network inference from scRNA data, popular in the community ^[45-47]^ and has shown strong performance in a recent benchmarking study ^[48]^. SCENIC’s three major steps are: First, it identifies groups of co-expressed genes as potential targets of a transcription factor (TF). Second, it filters these groups of genes to retain only TF targets with the corresponding binding motif, forming “regulons.” Third, it uses the AUCell method (embedded within SCENIC) to quantify the activity of each regulon in every cell. AUCell calculates the Area Under the Curve (AUC) for the regulon’s genes set in a ranking of all genes by expression for each cell. The top 20 regulons for each age groups were selected based on the regulon specificity score (RSS) (**Supplementary Figure 4B**). Unsupervised clustering on the Z score normalized regulon activity score matrix revealed clear differences in the GRN across different age groups (**Supplementary Figure 4C**). We hypothesize that the differences in the GRN might reflect differences in the pre- or postnatal developmental origins of the disease. Additional testing of GRN inference from individual studies shows that the high number of cells refines the overall GRN (**see Methods; Supplementary Figure 4D-E**).

To define gene regulatory programs (co-occurring gene modules, defined by a transcription factor and its targets) which are specific to different age groups (termed ‘regulon signature’), we used the clustered dendrogram to select the regulon clusters most associated with the pediatric (below 10 years-old) and adult samples (over 18 years-old) (**Figure3D; Supplementary Figure 4C**). The pediatric regulon signature, proposed to represent *in-utero* origin t(8;21) AML (henceforth termed ‘inferred-prenatal’), includes 16 regulons defined by a distinct group of hematopoietic transcription factors (TFs) (TRIM28, CTCF, RAD21, SOX4, TAL1, MYB, FOXN3, JUND, BCLAF1, ZBTB7A, IKZF1, MAZ, REST, YY1, CUX1, KDM5A), many of which have clearly defined roles in HSPCs and AML ^[49-51]^.The adult regulon signature, presumed representative of the postnatally acquired t(8;21) AML (henceforth termed ‘inferred-postnatal), combines 3 discrete clusters of regulons (YBX1, ENO1, and HDAC2; GATA1, POLE3, TFDP1, MYBL2, E2F4, and KLF1; IRF1, STAT1, IRF7, MAFF, ATF4, TAGLN2, SPI1, and KLF2), defined by TFs previously implicated in various hematopoietic, leukemic and inflammatory processes ^[52-54]^. Importantly, both signatures contain key components of the AP-1 complex, which is heavily implicated in the biology of t(8;21) AML ^[55, 56]^ and undergoes dynamic changes during aging ^[57]^. Samples of 6 individuals aged 12-17 clustered with the pediatric samples and showed enrichment for the inferred-prenatal signature (**Figure 3D**), suggesting that older adolescents (up to aged 17 in our cohort) more closely resemble pediatric AML with t(8;21) and remain biologically distinct from adult-onset disease. This implies that the inferred *in-utero* origin of t(8;21) AML can also be present in AML diagnosed in older children.

### Validation of Age-Associated Regulons in Bulk-RNA-Seq Cohorts of t(8;21) AML

We next sought to externally validate our age-associated regulon signatures in a larger cohort of patients. Bulk RNA-seq samples were obtained from the TARGET ^[2]^ and BeatAML ^[23]^ cohorts, selecting bone marrow samples taken at diagnosis in line with AML scAtlas data (n=83; **Supplementary Table 4**). We applied the AUCell algorithm from pySCENIC ^[22]^ to calculate the activity of our pediatric inferred-prenatal and adult inferred-postnatal regulons in each sample. Unsupervised clustering of the bulk RNA-seq AUCell results revealed discrete clusters of samples that were highly enriched for our inferred-prenatal and inferred-postnatal origin-associated regulons (**Figure 4A**).

Given the limitations of most scRNA-seq platforms in detecting lowly expressed genes, notably TFs, we leveraged bulk RNA-seq samples to refine our identified gene regulatory networks by detecting differentially expressed regulon-associated TFs. We used our inferred-prenatal and inferred-postnatal signature clusters and performed differential gene expression analysis between these samples, using two widely used tools (DESeq2^[59]^ and edgeR^[58]^) to ensure robustness of the results (**Figure 4B; Supplementary Figure 5A**). We then compared differentially expressed regulon-associated TFs between the two groups and intersected this with the differential genes detected by each method. Although changes in TF expression are subtle (**Figure 4B; Supplementary Figure 5A**), we identify significantly differentially expressed TFs which reflect the observed differences in regulon activity and indicate the most critical regulons in our age-related GRN signatures (**Figure 4A; Supplementary Figure 5B**). This further delineated the inferred-prenatal signature to 5 key TFs (KDM5A, REST, BCLAF1, YY1, and RAD21), and the inferred-postnatal signature to 8 TFs (ENO1, TFDP1, MYBL2, TAGLN2, KLF2, IRF7, SPI1, and YBX1).

**Figure 5.**
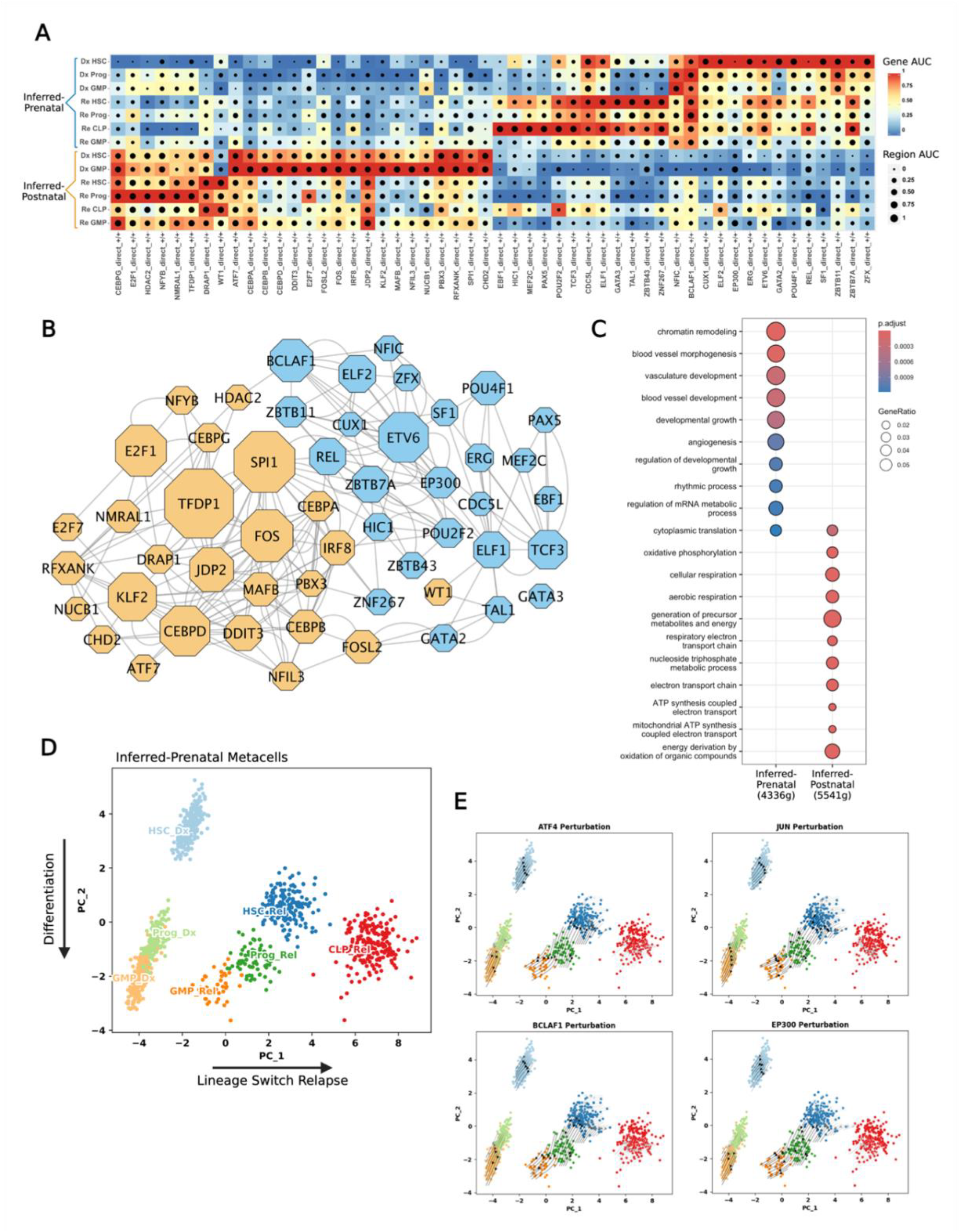
Combining Multiomics Data Interrogates Age-Associated Regulons. **(A)** SCENIC+ eRegulon dotplot of showing correlation between scRNA-seq target gene activity (indicated by the color scale) and scATAC-seq target region accessibility (depicted by spot size). RSS identified the key activating eRegulons (+/+) between inferred-prenatal and inferred-postnatal origin disease and allows comparison of diagnosis (Dx) and relapse (Rel) time points. (**B**) Network showing the inferred-prenatal (blue) and inferred-postnatal (orange) associated eRegulons. Node size represents the number of target genes in each regulon. Edges represent interactions between nodes. (**C**) Over-representation analysis of age-associated eRegulon target genes using GO Biological Processes curated gene sets. Adjusted P value threshold 0.05. (**D**) Principal components analysis (PCA) of the gene based eRegulon enrichment scores for the inferred-prenatal origin disease at diagnosis and relapse. PC1 axis explains variance occurring between diagnosis and relapse, where this patient underwent a lineage switch. PC2 captures variance related to hematopoietic differentiation. (**E**) SCENIC+ perturbation simulation shows the predicted effect of knockout of selected TFs on the previously computed PCA embedding. Arrows indicate the predicted shift in cell states relative to the initial PCA embedding.

We next performed gene set enrichment analysis (GSEA) on significantly differentially expressed genes as determined by edgeR^[58]^, to investigate pathways enriched in the inferred-prenatal samples compared to the inferred-postnatal ones (**Figure 4C**). Notably, inferred-prenatal samples showed increased expression of stemness-associated genes, and SMARCA2 target genes, a key player in HSC gene expression regulation and chromatin remodeling ^[60]^. SMARCA2 is also known to be upregulated during the fetal-to-adult HSC transition ^[61]^, implying that the observed SMARCA2 enrichment may indeed reflect the inferred fetal HSC cell-of-origin. Genes impacted by YY1 depletion were also downregulated compared to the samples of inferred-postnatal leukemia origin, which supports the identification of YY1 as an inferred-prenatal regulon (**Figure 4C**). To explore therapeutic implications, we performed GSEA using drug response signatures from published studies ^[62-65]^ (**Figure 4D**). This analysis revealed that inferred-prenatal origin t(8;21) AML is enriched for genes associated with increased chemosensitivity to cytarabine, venetoclax, and daunorubicin (**Figure 4D**).

We hypothesized that the increase in stemness-associated genes in the leukemia with the inferred-prenatal origin could be reflective of potential differences in the leukemic cell-of-origin and its impact on myeloid differentiation. We therefore performed cell type deconvolution using AutoGeneS ^[66]^, with a curated LSPC reference profile ^[7]^, to compare the cellular heterogeneity between prenatal and postnatal origin bulk RNA-seq samples (**Figure 4E**). This revealed a higher proportion of HSPC cell types (HSC, Prog), with a reduction in some differentiated myeloid cell types (ProMono-like, cDC-like) in the samples of inferred-prenatal origin (**Figure 4E**). To corroborate this finding, we examined cell type proportions in the original t(8;21) subset of AML scAtlas, confirming that cells with the inferred-prenatal signature comprise more HSCs than inferred-postnatal signature cells (**Supplementary Figure 5D-E**). However, comparison of cell type proportions in this dataset is confounded by differences in sample processing as some studies performed CD34 selection, hence there is more cell type diversity observed in the pediatric samples (**Supplementary Figure 5D-E**).

### Multiomics Single-Cell Data Reveals a Denoised GRN and Identifies Candidate Perturbations in Prenatal Origin t(8;21) AML

We next used the scRNA-seq and scATAC-seq data from a recent cohort of pediatric t(8;21) AML patients ^[24]^ at multiple clinical time points to uncover the enhancer-driven GRN (eGRN) in inferred-prenatal and inferred-postnatal origin t(8;21) AML (**Supplementary Table 4**). Initially, we identified two representative samples of our inferred-prenatal and inferred-postnatal signatures by using pySCENIC AUCell ^[22]^ to measure the activity of our previously defined regulons. Unsupervised clustering of the AUC scores was used to infer whether each sample matched the regulon signatures, identifying one inferred-prenatal sample and one inferred-postnatal sample for downstream analysis (**Supplementary Figure 6A**).

We then applied SCENIC+ ^[25]^, which integrates scRNA-seq and scATAC-seq to identify candidate enhancer regions and TF-binding motifs, linking TFs to target genes and identified enhancers. This creates enhancer-driven regulons (eRegulons), forming an eGRN. We applied SCENIC+ ^[25]^ to the leukemia samples with the inferred-prenatal and inferred-postnatal origin at diagnosis and relapse, keeping only regulons that showed a correlation between both modalities to retain only the most robust regulons (**Supplementary Figure 6B**). This revealed several eRegulons across both patients (**Supplementary Figure 6D**), many of which were patient specific, particularly when comparing HSPC populations (**Figure 4A**). The inferred-prenatal sample displayed a specific HSC eRegulon profile. In contrast, the inferred-postnatal sample more closely resembled the corresponding Granulocyte-Monocyte Progenitor (GMP) (**Figure 5A**). Interestingly, at relapse the inferred-prenatal origin patient undergoes a chemotherapy-driven lineage switch to a lymphoid phenotype, which may suggest that the leukemia originated from a less committed progenitor (**Figure 5A**).

To identify clusters of closely related eRegulons, we computed the correlations between eRegulons enrichment. We identified 2 main clusters of eRegulons which correspond to different inferred-signature samples (**Figure 4B; Supplementary Figure 6C**). For each eRegulon cluster, we used the associated target genes as input for gene ontology over representation analysis (ORA), to assess functional differences in the eGRN. This revealed fundamental differences in the underlying biological processes (**Figure 5C**; **Supplementary Figure 7A**). The AML sample with inferred-prenatal origin was enriched for many processes associated with development. In contrast, inferred-postnatal samples appeared more metabolism focused (**Figure 5C**; **Supplementary Figure 7A**). This further supports the association of these eRegulons with presumed prenatal origin t(8;21) AML, compared to postnatal origin disease.

Previous analysis using the TARGET ^[2]^ and BeatAML ^[23]^ datasets indicated that inferred-prenatal and inferred-postnatal origin t(8;21) AML may harbor different levels of chemosensitivity based on published drug response signatures (**Figure 4D**). Therefore, we performed *in silico* perturbations of eRegulon-associated TFs. PCA of the diagnosis and relapse samples recapitulated the expected differentiation trajectories along PC2, while separating diagnosis from relapse along PC1 (**Figure 5D**). Using the SCENIC+ ^[25]^ perturbation simulation workflow, we identified TFs estimated to induce differentiation, as defined by a negative shift in PC2 (**Supplementary Figure 7B**). We prioritized TFs predicted to impact the HSC compartment and identified 18 TFs with predicted significant effects on HSC differentiation (**Supplementary Figure 7C**). Several of these are components of the AP-1 complex (JUN, ATF4, FOSL2), which are established downstream targets of the t(8;21) fusion protein and are known to propagate t(8;21) AML ^[55, 67]^ (**Figure 5E; Supplementary Figure 7C**).

Using AP-1 complex members as a comparative baseline, we identified EP300 as one of the most impactful hits. EP300 has recently been shown to drive t(8;21) AML self-renewal through acetylation dependent mechanism ^[68]^. This suggests that presumed prenatal origin pediatric t(8;21) AML may be particularly sensitive to EP300 inhibition. One of the most striking predictions, for both diagnostic and relapse HSC populations, was BCLAF1 (**Figure 5E; Supplementary Figure 7C**). BCLAF1 is a regulator of normal HSPCs ^[69]^, and its expression level declines during hematopoietic differentiation. While recent studies have identified a role for BCLAF1 in AML ^[70]^, this has not been explored in detail in the context of pediatric AML or t(8;21) AML and may present a therapeutic opportunity.

We also performed SCENIC+ ^[25]^ perturbation modelling on the postnatal origin sample (AML12). In this case, the PCA was less straightforward to interpret, as branching differentiation trajectories towards a lymphoid or myeloid fate appear along the PC2 axis, while PC1 distinguishes diagnosis and relapse samples (**Supplementary Figure 7D**). Therefore, we prioritized TFs based on a predicted effect similar to AP-1 complex components, as it is known that this complex is a critical regulator in t(8;21) AML. We identified several TFs from our original postnatal origin signature were predicted to have an effect (**Supplementary Figures 7D-F**), supporting the relevance of the GRNs identified in our previous analyses.

To further investigate EP300 and BCLAF1, we queried the DepMap ^[71]^ database to assess the dependency of t(8;21) AML cell lines to these genes (**Supplementary Figure 7G**). We found that the two widely used cell lines of t(8;21) AML, KASUMI-1 (7-year-old donor) ^[72]^ and SKNO-1 (22-year-old donor) ^[73]^, were among the most sensitive to these perturbations based on their DepMap effect scores (**Supplementary Figure 7G**). Several other cell lines sensitive to BCLAF1 were derived from pediatric cancers, most notably neuroblastomas, which also arise *in-utero* ^[74]^ (**Supplementary Figure 7H**). Together, these findings suggest that EP300 inhibition may be particularly effective in t(8;21) AML, and that BCLAF1 may present a new therapeutic target for t(8;21) AML, particularly in pediatric cases with inferred pre-natal origin of the driver translocation.

## Discussion

Here we have generated a new data resource, AML scAtlas, to investigate AML biology across a broad range of subtypes at single-cell resolution. By including 222 samples comprising 748,679 cells of patients with a wide range of clinical characteristics, AML scAtlas overcomes the limitations of many standalone single-cell studies enabling AML subtype-focused analysis with enough data for robust statistical comparisons. This dataset is publicly available (https://cellxgene.bmh.manchester.ac.uk/AML/) providing the AML research community with a resource to address diverse biological questions and generate new hypotheses.

To further address a clinically relevant question using this data source, we compared differences between pediatric and adult-onset disease based on the potential biological effect of the *in-utero* origin of pediatric leukemia. Data of our AML scAtlas was used to explore the GRNs in adult and pediatric t(8;21) AML and revealed a strong age-associated GRN signature. This suggests that while pediatric and adult t(8;21) AMLs are propagated by the same driver translocation, they exhibit clear biological differences correlated with age. This may be due to differences in the cell-of-origin, with mouse models showing that t(8;21) AML can arise from a HSC or a more lineage restricted GMP ^[75]^. As pediatric t(8;21) can arise *in-utero*, as evidenced by previous studies ^[17]^, and adult t(8;21) is acquired postnatally ^[18, 19]^, we propose that the observed age-related differences in AML with t(8;21) reflect these differences in the developmental origins of the disease. We identified two distinct groups of regulons corresponding to either inferred-prenatal origin and inferred-postnatal origin disease. These regulons constitute the GRN underlying the cellular state, which can be informative when identifying molecular vulnerabilities to target leukemia.

Our cohort is the largest scRNA-seq dataset to explore t(8;21) AML biology to date, however, the number of patients included remains low (n=22), and many of the studies containing the adult samples used CD34 selection in their experimental protocol creating a bias towards HSPCs in these samples. To overcome some of these limitations, we used bulk RNA-seq samples from the TARGET ^[2]^ and BeatAML ^[23]^ studies with t(8;21) AML (n=83) to validate our regulon signatures. This identifies two clusters of samples which closely match these signatures, showing that the regulon patterns identified from our AML scAtlas are recapitulated with bulk RNA-seq data enabling exploration of larger patient cohorts. Comparisons between inferred-prenatal and inferred-postnatal origin transcriptomes prioritized TFs which were differentially expressed and highlighted differences in underlying biology and drug response. We identified 5 signature TFs (KDM5A, REST, BCLAF1, YY1, RAD21) for inferred-prenatal origin disease, several of which have roles in embryonic stem cells ^[76, 77]^, and critical functions in the maintenance of HSCs ^[49-51]^. In contrast, TFs identified in inferred-postnatal origin samples, such as interferon regulatory factors (IRFs), HDAC2, and SPI1, reflect inflammatory and immune processes, many of which have been implicated in leukemia ^[52-54]^. We also found that inferred-prenatal origin samples had a higher proportion of HSC/Prog cell types compared to inferred-postnatal origin samples, a more primitive state than postnatal onset t(8;21) AML cases, supporting the hypothesis that age-associated differences in the cell-of-origin influence disease biology.

Given these biological differences, we used bulk RNA-seq to predict chemosensitivity using published drug response signatures ^[62-65]^. Inferred-prenatal samples were enriched for genes indicative of cytarabine sensitivity and depleted of genes suggestive of daunorubicin and venetoclax resistance. These findings suggest that the developmental origins of the disease may influence drug responses, with potential implications in the design of novel therapeutic strategies and providing further biological evidence that pediatric AML might benefit from different clinical management compared with adult-onset AML. Importantly, venetoclax is currently in the AML23 trial (NCT05955261); our results support further evaluation of venetoclax treatment in pediatric t(8;21) AML.

Using an additional single-cell multiomic dataset, using SCENIC+, we reconstructed the eGRN in samples matching our inferred-prenatal and inferred-postnatal regulon signatures. Upon comparing eRegulons for each patient at diagnosis and relapse, we identified clusters of highly correlated eRegulons defined by different biological processes. Inferred-prenatal origin samples are characterized by developmental and transcriptional dysregulation, whereas inferred-postnatal origin samples are largely driven by fundamental cellular processes linked to inflammation. We used SCENIC+ to model the predicted impact of TF perturbations on our prenatal origin sample at diagnosis and relapse identified several key components of the AP-1 complex, which are critical in t(8;21) AML biology and are also associated with dynamic age-related transcriptional changes ^[55-57]^.

Through our analysis, we identified EP300 as a candidate target, which has been shown to be critical for t(8;21) AML biology ^[68]^ with demonstrable effects in KASUMI-1 and SKNO1 cell lines. EP300 has been identified as a promising therapeutic target in AML with several molecules in development ^[78]^; our data indicate potential specific therapeutic benefit in prenatal origin t(8;21) AML. One of the most impactful perturbation predictions for the HSC compartment at diagnosis and relapse was BCLAF1. This is consistent with previous evidence of its importance in HSCs ^[69, 79]^ and AML ^[70]^, but has not been studied specifically in the context of pediatric AML or t(8;21) AML previously. The DepMap data shows that KASUMI-1 is the most sensitive myeloid cell line to BCLAF1 perturbation, and our GRN analyses suggest it is particularly active in pediatric t(8;21) AML of inferred *in-utero* origin, thus this may represent an additional prognostic indicator.

Further investigations are required to characterize the roles of both EP300 and BCLAF1 in prenatal origin t(8;21) AML before any clinical realisation. EP300 has already been investigated as a target in AML, so future work should focus on the pediatric AML setting with *in-vitro* and *in-vivo* studies using EP300/CBP inhibitors such as inobrodib ^[78]^. In contrast

BCLAF1 is relatively unexplored, and additional work is required to elucidate its molecular function and assess its potential as a therapeutic target. BCLAF1 may ultimately prove most valuable as a biomarker of *in-utero* t(8;21) AML, enabling distinction between late-onset *in-utero* and postnatal disease. This would require molecular validation in a large cohort of pediatric patients with Gunthrie spots to confirm whether they had acquired t(8;21) *in-utero*.

## Conclusions

Overall, our study demonstrates that large-scale single-cell data integration is a powerful approach to dissect specific patient groups in detail, and enabling robust comparative analyses. We present the AML scAtlas as a publicly available resource for the research community to address diverse biological questions. By applying AML scAtlas to t(8;21) AML, we identified age-associated gene regulatory networks that likely reflect differences in the developmental origins, biology and outcome of the disease. These findings also highlight novel candidate therapeutic targets which may be more relevant in pediatric t(8;21) AML compared to adult-onset disease, offering opportunities for more tailored treatment strategies.

## Methods

For the complete analysis code, including the conda environments used for analysis, see GitHub Repo (https://github.com/jesswhitts/AML-scAtlas).

### Data Collection

A literature search was performed for published AML scRNA-seq datasets ^[4-6, 8, 26-40]^. Suitable studies were selected based on the data quality (over 1000 counts and 500 genes detected per cell for most of the data). Diagnostic, primary AML samples were selected from each AML study. Where healthy donor samples were present, these were also included, along with an additional 4 studies with healthy bone marrow samples.

### Initial Data Processing

Each scRNA-seq dataset underwent initial quality control individually using Scanpy (v1.9.3) ^[80]^ as some studies provided raw data and others provided pre-filtered data. Where raw data was provided, doublets were removed using Scrublet (v0.2.3) ^[81]^ and cells were filtered using the median absolute deviation as described in this single-cell best practices handbook ^[10, 82]^.

Once filtered, datasets were combined, and quality control was performed using Scanpy (v1.9.3) ^[80]^. The full dataset had quality thresholds applied (percentage mitochondrial counts <10, read counts >1000, gene counts >500), removing any samples which had fewer than 50 cells remaining after filtering. Genes present in <50 cells were removed. MATAL1 was removed as this was highly abundant in many cells and considered artefactual.

### Batch Correction

The presence of batch effects was determined through dimensionality reduction and clustering using Scanpy (v1.9.3) ^[80]^ and using the kBET algorithm (v0.99.6) ^[83]^. This was repeated on individual studies, to assess whether there were sample-wise batch effects. Batch correction benchmarking was implemented using Harmony (Scanpy v1.9.3 implementation) ^[11]^, scVI (v1.0.3) ^[12]^, and scANVI (v1.0.3) ^[13]^ and quantified using scIB (1.1.4) ^[9]^. Different numbers of highly variable genes were used to select the optimal number for integration. Batch correction was performed using scVI (v1.0.3) ^[12]^ with the top 2000 highly variable genes, using sample as the model covariate.

### AML scAtlas Cell Type Annotation

The scVI corrected embedding was used to run UMAP and Leiden clustering using Scanpy functions (v1.9.3) ^[80]^. Cell type annotation was performed using CellTypist (v1.6.0) ^[84]^ using the ‘Immune_All_Low.pkl’ model, SingleR (v2.0.0) ^[85]^ using the Novershtern hematopoietic refreference^[86]^, and scType (v1.0) ^[87]^ with the tissue defined as ‘Immune system’. Full automated tool outputs are detailed in **Supplementary Table 3**; overall we found that the results varied significantly between different tools. We postulate that this is, in part, due to differences in the reference profiles used. Thus, we opted to use the best consensus of these different tools for our cluster identity assignments.

### AML scAtlas LSC Annotation

HSPC clusters were selected from AML scAtlas, and the scVI corrected embedding was used to re-compute UMAP using Scanpy functions (v1.9.3) ^[80]^. As our previous cell type annotations used generic reference profiles and were not AML specific, we generated a custom cell type annotation reference to identify LSCs. We created a custom SingleR (v2.0.0) ^[85]^ reference using the Zeng et al ^[7]^ revised annotations of the Van Galen et al ^[5]^ dataset (**Supplementary Table 3**). This was also correlated with the LSC6 ^[43]^ and LSC17 ^[44]^ scores for each cell. To compare LSC abundance between ELN risk groups, chi2_contingency was implemented from SciPy (v1.12.0).

AML with t(8;21) Analysis

Samples with the t(8;21) translocation were selected from the full AML scAtlas. The UMAP was re-computed, and genes were filtered to remove those detected in fewer than 50 cells for the revised dataset, leaving 24,866 genes remaining. Gene regulatory network analysis was performed using pySCENIC ^[21, 22]^ (v0.12.1) as per the recommended workflow. To facilitate comparisons between age groups, cell types were focused on HSPCs, as many adult samples were originally enriched for CD34. The RSS was calculated for the adult and pediatric samples to select the top 20 differential regulons per age group. Using SciPy hierarchical clustering (v1.12.0), regulons were filtered to identify regulon signature groups used for downstream analysis.

### Bulk RNA-Seq Analysis

Bulk RNA-Seq data was downloaded for the TARGET ^[2]^ and BeatAML ^[23]^ cohorts and samples with t(8;21) were selected. Only bone marrow samples taken at diagnosis were used for downstream analyses (**Supplementary Table 4**). Using the previously defined signature regulons, the AUCell algorithm ^[21]^ (v0.12.1) was implemented to measure regulon activity. Hierarchical clustering was performed using SciPy v1.12.0) to identify samples most enriched for each age-related signature. Differential gene expression analysis was implemented using edgeR ^[58]^ (v3.42.4) and DESeq2 ^[59]^ (v1.40.2), using a log2 fold change threshold of 0.5 and an adjusted p value cutoff of 0.01. Candidate differential genes, ranked on log2 fold change, underwent GSEA with GSEApy (v0.10.8) using a significance threshold of 0.05. Cell type deconvolution was performed using AutoGeneS ^[66]^ (v1.0.4) using the recommended workflow. Significance when comparing groups was ascertained using a student’s T-test on the predicted cell type proportion values for each sample.

### Single-Cell Multi-Omics Analysis

The Lambo et al ^[24]^ scRNA-seq and scATAC-seq data from pediatric AML bone marrow samples was downloaded, and the t(8;21) samples selected (**Supplementary Table 4**). Using our previously defined signature regulons, the AUCell algorithm ^[21]^ (v0.12.1) was implemented to measure regulon activity. This identified the samples most enriched for each age-related signature as AML16 and AML12.

The SCENIC+ pipeline ^[25]^ (v1.0a1) was implemented as per the recommended Snakemake workflow for creating pseudo-multiome data. Regulons were filtered by correlation between modalities, using a threshold of 0.2 for non-multiome data. The most robust regulons were prioritized based on the SCENIC+ recommendations (direct +/+) ^[25]^. To facilitate comparisons, the eRegulon RSS was calculated for each patient and the top 30 eRegulons selected. The correlation between the gene sets underpinning eRegulons was calculated and sample-associated clusters were selected for over-representation analysis with clusterProfiler ^[88]^ (v4.8.3).

To predict the impact of specific TF perturbations on key cell types, SCENIC+ ^[25]^ perturbation modelling was implemented using the recommended parameters. TFs were then prioritized on their predicted impact on HSC differentiation and visualised using the PCA embedding. Candidate targets EP300 and BCLAF1 were queried in the DepMap ^[71]^ databases to infer their potential importance.

## Availability of Data and Materials

The AML scAtlas is hosted online for public use (https://cellxgene.bmh.manchester.ac.uk/AML/). The processed AnnData object is also available to download from figshare (DOI: 10.48420/27269946). Details of all data used in this study can be found in Supplementary Table 1, along with associated links to the original data. All code used to perform the analyses presented here can be accessed in the GitHub repository: (https://github.com/jesswhitts/AML-scAtlas). The samples used for validation analyses are publicly available and are detailed in Supplementary Table 4. The SCENIC+ eGRN files are provided as Supplementary Table 5.

## Author Contributions

MI, SMB, and GL conceived the study and oversaw the research. JW did the data collection and implemented the analyses. SM guided sub-analyses and assisted with interpretation. JW wrote the first draft of the manuscript. All authors interpreted the data and edited the manuscript. All authors approved the final manuscript.

## Acknowledgements

JW was funded by MRC DTP award (MR/W007428/1), SM by Blood cancer UK (15038) and CCLG (2016 09), while GL by Cancer Research UK (C5759/A20971 & C5759/A27412) and MI by MRC (MR/X014088/1).

We would like to thank all authors of the public data used in this study for their contributions to scientific community. We also acknowledge useful discussions around this work with Magnus Rattray.

## References

1. Tenen DG. Disruption of differentiation in human cancer: AML shows the way. Nature Reviews Cancer. 2003;3(2):89–101.

2. Bolouri H, Farrar JE, Triche T, Jr., Ries RE, Lim EL, Alonzo TA, Ma Y, Moore R, Mungall AJ, Marra MA, Zhang J, Ma X, Liu Y, Liu Y, Auvil JMG, Davidsen TM, Gesuwan P, Hermida LC, Salhia B, Capone S, Ramsingh G, Zwaan CM, Noort S, Piccolo SR, Kolb EA, Gamis AS, Smith MA, Gerhard DS, Meshinchi S. The molecular landscape of pediatric acute myeloid leukemia reveals recurrent structural alterations and age-specific mutational interactions. Nat Med. 2018;24(1):103–12.

3. Velten L, Haas SF, Raffel S, Blaszkiewicz S, Islam S, Hennig BP, Hirche C, Lutz C, Buss EC, Nowak D, Boch T, Hofmann WK, Ho AD, Huber W, Trumpp A, Essers MA, Steinmetz LM. Human haematopoietic stem cell lineage commitment is a continuous process. Nat Cell Biol. 2017;19(4):271–81.

4. Velten L, Story BA, Hernández-Malmierca P, Raffel S, Leonce DR, Milbank J, Paulsen M, Demir A, Szu-Tu C, Frömel R, Lutz C, Nowak D, Jann J-C, Pabst C, Boch T, Hofmann W-K, Müller-Tidow C, Trumpp A, Haas S, Steinmetz LM. Identification of leukemic and pre-leukemic stem cells by clonal tracking from single-cell transcriptomics. Nature Communications. 2021;12(1):1366.

5. van Galen P, Hovestadt V, Wadsworth Ii MH, Hughes TK, Griffin GK, Battaglia S, Verga JA, Stephansky J, Pastika TJ, Lombardi Story J, Pinkus GS, Pozdnyakova O, Galinsky I, Stone RM, Graubert TA, Shalek AK, Aster JC, Lane AA, Bernstein BE. Single-Cell RNA-Seq Reveals AML Hierarchies Relevant to Disease Progression and Immunity. Cell. 2019;176(6):1265-81.e24.

6. Beneyto-Calabuig S, Merbach AK, Kniffka JA, Antes M, Szu-Tu C, Rohde C, Waclawiczek A, Stelmach P, Gräßle S, Pervan P, Janssen M, Landry JJM, Benes V, Jauch A, Brough M, Bauer M, Besenbeck B, Felden J, Bäumer S, Hundemer M, Sauer T, Pabst C, Wickenhauser C, Angenendt L, Schliemann C, Trumpp A, Haas S, Scherer M, Raffel S, Müller-Tidow C, Velten L. Clonally resolved single-cell multi-omics identifies routes of cellular differentiation in acute myeloid leukemia. Cell Stem Cell. 2023;30(5):706-21.e8.

7. Zeng AGX, Bansal S, Jin L, Mitchell A, Chen WC, Abbas HA, Chan-Seng-Yue M, Voisin V, van Galen P, Tierens A, Cheok M, Preudhomme C, Dombret H, Daver N, Futreal PA, Minden MD, Kennedy JA, Wang JCY, Dick JE. A cellular hierarchy framework for understanding heterogeneity and predicting drug response in acute myeloid leukemia. Nature Medicine. 2022;28(6):1212–23.

8. Stetson LC, Balasubramanian D, Ribeiro SP, Stefan T, Gupta K, Xu X, Fourati S, Roe A, Jackson Z, Schauner R, Sharma A, Tamilselvan B, Li S, de Lima M, Hwang TH, Balderas R, Saunthararajah Y, Maciejewski J, LaFramboise T, Barnholtz-Sloan JS, Sekaly RP, Wald DN. Single cell RNA sequencing of AML initiating cells reveals RNA-based evolution during disease progression. Leukemia. 2021;35(10):2799–812.

9. Luecken MD, Büttner M, Chaichoompu K, Danese A, Interlandi M, Mueller MF, Strobl DC, Zappia L, Dugas M, Colomé-Tatché M, Theis FJ. Benchmarking atlas-level data integration in single-cell genomics. Nature Methods. 2022;19(1):41–50.

10. Heumos L, Schaar AC, Lance C, Litinetskaya A, Drost F, Zappia L, Lücken MD, Strobl DC, Henao J, Curion F, Aliee H, Ansari M, Badia-i-Mompel P, Büttner M, Dann E, Dimitrov D, Dony L, Frishberg A, He D, Hediyeh-zadeh S, Hetzel L, Ibarra IL, Jones MG, Lotfollahi M, Martens LD, Müller CL, Nitzan M, Ostner J, Palla G, Patro R, Piran Z, Ramírez-Suástegui C, Saez-Rodriguez J, Sarkar H, Schubert B, Sikkema L, Srivastava A, Tanevski J, Virshup I, Weiler P, Schiller HB, Theis FJ, Single-cell Best Practices C. Best practices for single-cell analysis across modalities. Nature Reviews Genetics. 2023.

11. Korsunsky I, Millard N, Fan J, Slowikowski K, Zhang F, Wei K, Baglaenko Y, Brenner M, Loh P-r, Raychaudhuri S. Fast, sensitive and accurate integration of single-cell data with Harmony. Nature Methods. 2019;16(12):1289–96.

12. Lopez R, Regier J, Cole MB, Jordan MI, Yosef N. Deep generative modeling for single-cell transcriptomics. Nature Methods. 2018;15(12):1053–8.

13. Xu C, Lopez R, Mehlman E, Regier J, Jordan MI, Yosef N. Probabilistic harmonization and annotation of single-cell transcriptomics data with deep generative models. Molecular Systems Biology. 2021;17(1):e9620.

14. Balgobind BV, Van den Heuvel-Eibrink MM, De Menezes RX, Reinhardt D, Hollink IH, Arentsen-Peters ST, van Wering ER, Kaspers GJ, Cloos J, de Bont ES, Cayuela JM, Baruchel A, Meyer C, Marschalek R, Trka J, Stary J, Beverloo HB, Pieters R, Zwaan CM, den Boer ML. Evaluation of gene expression signatures predictive of cytogenetic and molecular subtypes of pediatric acute myeloid leukemia. Haematologica. 2011;96(2):221–30.

15. Wiggers CRM, Baak ML, Sonneveld E, Nieuwenhuis EES, Bartels M, Creyghton MP. AML Subtype Is a Major Determinant of the Association between Prognostic Gene Expression Signatures and Their Clinical Significance. Cell Reports. 2019;28(11):2866-77.e5.

16. Chaudhury S, O’Connor C, Cañete A, Bittencourt-Silvestre J, Sarrou E, Prendergast Á, Choi J, Johnston P, Wells CA, Gibson B, Keeshan K. Age-specific biological and molecular profiling distinguishes paediatric from adult acute myeloid leukaemias. Nat Commun. 2018;9(1):5280.

17. Wiemels JL, Xiao Z, Buffler PA, Maia AT, Ma X, Dicks BM, Smith MT, Zhang L, Feusner J, Wiencke J, Pritchard-Jones K, Kempski H, Greaves M. In utero origin of t(8;21) AML1-ETO translocations in childhood acute myeloid leukemia. Blood. 2002;99(10):3801–5.

18. Welch JS, Ley TJ, Link DC, Miller CA, Larson DE, Koboldt DC, Wartman LD, Lamprecht TL, Liu F, Xia J, Kandoth C, Fulton RS, McLellan MD, Dooling DJ, Wallis JW, Chen K, Harris CC, Schmidt HK, Kalicki-Veizer JM, Lu C, Zhang Q, Lin L, O’Laughlin MD, McMichael JF, Delehaunty KD, Fulton LA, Magrini VJ, McGrath SD, Demeter RT, Vickery TL, Hundal J, Cook LL, Swift GW, Reed JP, Alldredge PA, Wylie TN, Walker JR, Watson MA, Heath SE, Shannon WD, Varghese N, Nagarajan R, Payton JE, Baty JD, Kulkarni S, Klco JM, Tomasson MH, Westervelt P, Walter MJ, Graubert TA, DiPersio JF, Ding L, Mardis ER, Wilson RK. The origin and evolution of mutations in acute myeloid leukemia. Cell. 2012;150(2):264–78.

19. Jaiswal S, Fontanillas P, Flannick J, Manning A, Grauman PV, Mar BG, Lindsley RC, Mermel CH, Burtt N, Chavez A, Higgins JM, Moltchanov V, Kuo FC, Kluk MJ, Henderson B, Kinnunen L, Koistinen HA, Ladenvall C, Getz G, Correa A, Banahan BF, Gabriel S, Kathiresan S, Stringham HM, McCarthy MI, Boehnke M, Tuomilehto J, Haiman C, Groop L, Atzmon G, Wilson JG, Neuberg D, Altshuler D, Ebert BL. Age-related clonal hematopoiesis associated with adverse outcomes. N Engl J Med. 2014;371(26):2488–98.

20. National Cancer Registration and Analysis Service, Northern Ireland Cancer Registry, Scottish Cancer Registry, Unit WCIaS. Children, teenagers and young adults UK cancer statistics report 2021 2021 [Available from: http://www.ncin.org.uk/cancer_type_and_topic_specific_work/cancer_type_specific_work/cancer_in_children_teenagers_and_young_adults/.

21. Aibar S, González-Blas Cb, Moerman T, Huynh-Thu VA, Imrichova H, Hulselmans G, Rambow F, Marine J-C, Geurts P, Aerts J, van den Oord J, Atak ZK, Wouters J, Aerts S. SCENIC: single-cell regulatory network inference and clustering. Nature Methods. 2017;14(11):1083–6.

22. Van de Sande B, Flerin C, Davie K, De Waegeneer M, Hulselmans G, Aibar S, Seurinck R, Saelens W, Cannoodt R, Rouchon Q, Verbeiren T, De Maeyer D, Reumers J, Saeys Y, Aerts S. A scalable SCENIC workflow for single-cell gene regulatory network analysis. Nature Protocols. 2020;15(7):2247–76.

23. Burd A, Levine RL, Ruppert AS, Mims AS, Borate U, Stein EM, Patel P, Baer MR, Stock W, Deininger M, Blum W, Schiller G, Olin R, Litzow M, Foran J, Lin TL, Ball B, Boyiadzis M, Traer E, Odenike O, Arellano M, Walker A, Duong VH, Kovacsovics T, Collins R, Shoben AB, Heerema NA, Foster MC, Vergilio J-A, Brennan T, Vietz C, Severson E, Miller M, Rosenberg L, Marcus S, Yocum A, Chen T, Stefanos M, Druker B, Byrd JC. Precision medicine treatment in acute myeloid leukemia using prospective genomic profiling: feasibility and preliminary efficacy of the Beat AML Master Trial. Nature Medicine. 2020;26(12):1852–8.

24. Lambo S, Trinh DL, Ries RE, Jin D, Setiadi A, Ng M, Leblanc VG, Loken MR, Brodersen LE, Dai F, Pardo LM, Ma X, Vercauteren SM, Meshinchi S, Marra MA. A longitudinal single-cell atlas of treatment response in pediatric AML. Cancer Cell. 2023.

25. Bravo González-Blas C, De Winter S, Hulselmans G, Hecker N, Matetovici I, Christiaens V, Poovathingal S, Wouters J, Aibar S, Aerts S. SCENIC+: single-cell multiomic inference of enhancers and gene regulatory networks. Nature Methods. 2023;20(9):1355–67.

26. Zheng GXY, Terry JM, Belgrader P, Ryvkin P, Bent ZW, Wilson R, Ziraldo SB, Wheeler TD, McDermott GP, Zhu J, Gregory MT, Shuga J, Montesclaros L, Underwood JG, Masquelier DA, Nishimura SY, Schnall-Levin M, Wyatt PW, Hindson CM, Bharadwaj R, Wong A, Ness KD, Beppu LW, Deeg HJ, McFarland C, Loeb KR, Valente WJ, Ericson NG, Stevens EA, Radich JP, Mikkelsen TS, Hindson BJ, Bielas JH. Massively parallel digital transcriptional profiling of single cells. Nature Communications. 2017;8(1):14049.

27. Petti AA, Williams SR, Miller CA, Fiddes IT, Srivatsan SN, Chen DY, Fronick CC, Fulton RS, Church DM, Ley TJ. A general approach for detecting expressed mutations in AML cells using single cell RNA-sequencing. Nat Commun. 2019;10(1):3660.

28. Jiang L, Li XP, Dai YT, Chen B, Weng XQ, Xiong SM, Zhang M, Huang JY, Chen Z, Chen SJ. Multidimensional study of the heterogeneity of leukemia cells in t(8;21) acute myelogenous leukemia identifies the subtype with poor outcome. Proc Natl Acad Sci U S A. 2020;117(33):20117–26.

29. Johnston G, Ramsey HE, Liu Q, Wang J, Stengel KR, Sampathi S, Acharya P, Arrate M, Stubbs MC, Burn T, Savona MR, Hiebert SW. Nascent transcript and single-cell RNA-seq analysis defines the mechanism of action of the LSD1 inhibitor INCB059872 in myeloid leukemia. Gene. 2020;752:144758.

30. Pei S, Pollyea DA, Gustafson A, Stevens BM, Minhajuddin M, Fu R, Riemondy KA, Gillen AE, Sheridan RM, Kim J, Costello JC, Amaya ML, Inguva A, Winters A, Ye H, Krug A, Jones CL, Adane B, Khan N, Ponder J, Schowinsky J, Abbott D, Hammes A, Myers JR, Ashton JM, Nemkov T, D’Alessandro A, Gutman JA, Ramsey HE, Savona MR, Smith CA, Jordan CT. Monocytic Subclones Confer Resistance to Venetoclax-Based Therapy in Patients with Acute Myeloid Leukemia. Cancer Discov. 2020;10(4):536–51.

31. Li K, Du Y, Cai Y, Liu W, Lv Y, Huang B, Zhang L, Wang Z, Liu P, Sun Q, Li N, Zhu M, Bosco B, Li L, Wu W, Wu L, Li J, Wang Q, Hong M, Qian S. Single-cell analysis reveals the chemotherapy-induced cellular reprogramming and novel therapeutic targets in relapsed/refractory acute myeloid leukemia. Leukemia. 2023;37(2):308–25.

32. Lasry A, Nadorp B, Fornerod M, Nicolet D, Wu H, Walker CJ, Sun Z, Witkowski MT, Tikhonova AN, Guillamot-Ruano M, Cayanan G, Yeaton A, Robbins G, Obeng EA, Tsirigos A, Stone RM, Byrd JC, Pounds S, Carroll WL, Gruber TA, Eisfeld A-K, Aifantis I. An inflammatory state remodels the immune microenvironment and improves risk stratification in acute myeloid leukemia. Nature Cancer. 2023;4(1):27–42.

33. Fiskus W, Mill CP, Birdwell C, Davis JA, Das K, Boettcher S, Kadia TM, DiNardo CD, Takahashi K, Loghavi S, Soth MJ, Heffernan T, McGeehan GM, Ruan X, Su X, Vakoc CR, Daver N, Bhalla KN. Targeting of epigenetic co-dependencies enhances anti-AML efficacy of Menin inhibitor in AML with MLL1-r or mutant NPM1. Blood Cancer Journal. 2023;13(1):53.

34. Naldini MM, Casirati G, Barcella M, Rancoita PMV, Cosentino A, Caserta C, Pavesi F, Zonari E, Desantis G, Gilioli D, Carrabba MG, Vago L, Bernardi M, Di Micco R, Di Serio C, Merelli I, Volpin M, Montini E, Ciceri F, Gentner B. Longitudinal single-cell profiling of chemotherapy response in acute myeloid leukemia. Nature Communications. 2023;14(1):1285.

35. Mumme H, Thomas BE, Bhasin SS, Krishnan U, Dwivedi B, Perumalla P, Sarkar D, Ulukaya GB, Sabnis HS, Park SI, DeRyckere D, Raikar SS, Pauly M, Summers RJ, Castellino SM, Wechsler DS, Porter CC, Graham DK, Bhasin M. Single-cell analysis reveals altered tumor microenvironments of relapse- and remission-associated pediatric acute myeloid leukemia. Nature Communications. 2023;14(1):6209.

36. Zhang Y, Jiang S, He F, Tian Y, Hu H, Gao L, Zhang L, Chen A, Hu Y, Fan L, Yang C, Zhou B, Liu D, Zhou Z, Su Y, Qin L, Wang Y, He H, Lu J, Xiao P, Hu S, Wang Q-F. Single-cell transcriptomics reveals multiple chemoresistant properties in leukemic stem and progenitor cells in pediatric AML. Genome Biology. 2023;24(1):199.

37. Li B, Kowalczyk MS, Slyper M, Jellert G, Tabaka M, Ashenberg O, Waldman J, Dionne D, Abigail K, Hui M, Yang Y, Rozenblatt-Rosen O, Regev A. A single cell immune cell atlas of human hematopoietic system Human Cell Atlas Data Portal: Human Cell Atlas; 2022 [Available from: https://explore.data.humancellatlas.org/projects/cc95ff89-2e68-4a08-a234-480eca21ce79.

38. Oetjen KA, Lindblad KE, Goswami M, Gui G, Dagur PK, Lai C, Dillon LW, McCoy JP, Hourigan CS. Human bone marrow assessment by single-cell RNA sequencing, mass cytometry, and flow cytometry. JCI Insight. 2018;3(23).

39. Setty M, Kiseliovas V, Levine J, Gayoso A, Mazutis L, Pe’er D. Characterization of cell fate probabilities in single-cell data with Palantir. Nat Biotechnol. 2019;37(4):451–60.

40. Caron M, St-Onge P, Sontag T, Wang YC, Richer C, Ragoussis I, Sinnett D, Bourque G. Single-cell analysis of childhood leukemia reveals a link between developmental states and ribosomal protein expression as a source of intra-individual heterogeneity. Scientific Reports. 2020;10(1):8079.

41. Döhner H, Wei AH, Appelbaum FR, Craddock C, DiNardo CD, Dombret H, Ebert BL, Fenaux P, Godley LA, Hasserjian RP, Larson RA, Levine RL, Miyazaki Y, Niederwieser D, Ossenkoppele G, Röllig C, Sierra J, Stein EM, Tallman MS, Tien H-F, Wang J, Wierzbowska A, Löwenberg B. Diagnosis and management of AML in adults: 2022 recommendations from an international expert panel on behalf of the ELN. Blood. 2022;140(12):1345–77.

42. Montefiori LE, Bendig S, Gu Z, Chen X, Pölönen P, Ma X, Murison A, Zeng A, Garcia-Prat L, Dickerson K, Iacobucci I, Abdelhamed S, Hiltenbrand R, Mead PE, Mehr CM, Xu B, Cheng Z, Chang TC, Westover T, Ma J, Stengel A, Kimura S, Qu C, Valentine MB, Rashkovan M, Luger S, Litzow MR, Rowe JM, den Boer ML, Wang V, Yin J, Kornblau SM, Hunger SP, Loh ML, Pui CH, Yang W, Crews KR, Roberts KG, Yang JJ, Relling MV, Evans WE, Stock W, Paietta EM, Ferrando AA, Zhang J, Kern W, Haferlach T, Wu G, Dick JE, Klco JM, Haferlach C, Mullighan CG. Enhancer Hijacking Drives Oncogenic BCL11B Expression in Lineage-Ambiguous Stem Cell Leukemia. Cancer Discov. 2021;11(11):2846–67.

43. Elsayed AH, Rafiee R, Cao X, Raimondi S, Downing JR, Ribeiro R, Fan Y, Gruber TA, Baker S, Klco J, Rubnitz JE, Pounds S, Lamba JK. A six-gene leukemic stem cell score identifies high risk pediatric acute myeloid leukemia. Leukemia. 2020;34(3):735–45.

44. Ng SWK, Mitchell A, Kennedy JA, Chen WC, McLeod J, Ibrahimova N, Arruda A, Popescu A, Gupta V, Schimmer AD, Schuh AC, Yee KW, Bullinger L, Herold T, Görlich D, Büchner T, Hiddemann W, Berdel WE, Wörmann B, Cheok M, Preudhomme C, Dombret H, Metzeler K, Buske C, Löwenberg B, Valk PJM, Zandstra PW, Minden MD, Dick JE, Wang JCY. A 17-gene stemness score for rapid determination of risk in acute leukaemia. Nature. 2016;540(7633):433–7.

45. Hamed AA, Kunz DJ, El-Hamamy I, Trinh QM, Subedar OD, Richards LM, Foltz W, Bullivant G, Ware M, Vladoiu MC, Zhang J, Raj AM, Pugh TJ, Taylor MD, Teichmann SA, Stein LD, Simons BD, Dirks PB. A brain precursor atlas reveals the acquisition of developmental-like states in adult cerebral tumours. Nat Commun. 2022 Jul 19;13(1):4178.

46. Barnett SN, Cujba AM, Yang L, Maceiras AR, Li S, Kedlian VR, Pett JP, Polanski K, Miranda AMA, Xu C, Cranley J, Kanemaru K, Lee M, Mach L, Perera S, Tudor C, Joseph PD, Pritchard S, Toscano-Rivalta R, Tuong ZK, Bolt L, Petryszak R, Prete M, Cakir B, Huseynov A, Sarropoulos I, Chowdhury RA, Elmentaite R, Madissoon E, Oliver AJ, Campos L, Brazovskaja A, Gomes T, Treutlein B, Kim CN, Nowakowski TJ, Meyer KB, Randi AM, Noseda M, Teichmann SA. An organotypic atlas of human vascular cells. Nat Med. 2024 Dec;30(12):3468–3481.

47. Zhang B, He P, Lawrence JEG, Wang S, Tuck E, Williams BA, Roberts K, Kleshchevnikov V, Mamanova L, Bolt L, Polanski K, Li T, Elmentaite R, Fasouli ES, Prete M, He X, Yayon N, Fu Y, Yang H, Liang C, Zhang H, Blain R, Chedotal A, FitzPatrick DR, Firth H, Dean A, Bayraktar OA, Marioni JC, Barker RA, Storer MA, Wold BJ, Zhang H, Teichmann SA. A human embryonic limb cell atlas resolved in space and time. Nature. 2024 Nov;635(8039):668–678.

48. Nguyen H, Tran D, Tran B, Pehlivan B, Nguyen T. A comprehensive survey of regulatory network inference methods using single cell RNA sequencing data. Brief Bioinform. 2021 May 20;22(3):bbaa190.

49. Lu Z, Hong CC, Kong G, Assumpção A, Ong IM, Bresnick EH, Zhang J, Pan X. Polycomb Group Protein YY1 Is an Essential Regulator of Hematopoietic Stem Cell Quiescence. Cell Rep. 2018;22(6):1545–59.

50. Fisher JB, Peterson J, Reimer M, Stelloh C, Pulakanti K, Gerbec ZJ, Abel AM, Strouse JM, Strouse C, McNulty M, Malarkannan S, Crispino JD, Milanovich S, Rao S. The cohesin subunit Rad21 is a negative regulator of hematopoietic self-renewal through epigenetic repression of Hoxa7 and Hoxa9. Leukemia. 2017;31(3):712–9.

51. Kumar P, Zhang N, Lee J, Cheng H, Kurtz K, Conneely SE, Sasidharan R, Rau RE, Pati D. Cohesin Subunit RAD21 Regulates the Differentiation and Self-Renewal of Hematopoietic Stem and Progenitor Cells. Stem Cells. 2023;41(10):971–85.

52. Ning S, Pagano JS, Barber GN. IRF7: activation, regulation, modification and function. Genes & Immunity. 2011;12(6):399–414.

53. Fischer J, Walter C, Tönges A, Aleth H, Jordão MJC, Leddin M, Gröning V, Erdmann T, Lenz G, Roth J, Vogl T, Prinz M, Dugas M, Jacobsen ID, Rosenbauer F. Safeguard function of PU.1 shapes the inflammatory epigenome of neutrophils. Nature Immunology. 2019;20(5):546–58.

54. Fang W-F, Chen Y-M, Lin C-Y, Huang H-L, Yeh H, Chang Y-T, Huang K-T, Lin M-C. Histone deacetylase 2 (HDAC2) attenuates lipopolysaccharide (LPS)-induced inflammation by regulating PAI-1 expression. Journal of Inflammation. 2018;15(1):3.

55. Ptasinska A, Pickin A, Assi SA, Chin PS, Ames L, Avellino R, Gröschel S, Delwel R, Cockerill PN, Osborne CS, Bonifer C. RUNX1-ETO Depletion in t(8;21) AML Leads to C/EBPα- and AP-1-Mediated Alterations in Enhancer-Promoter Interaction. Cell Reports. 2019;28(12):3022-31.e7.

56. Martinez-Soria N, McKenzie L, Draper J, Ptasinska A, Issa H, Potluri S, Blair HJ, Pickin A, Isa A, Chin PS, Tirtakusuma R, Coleman D, Nakjang S, Assi S, Forster V, Reza M, Law E, Berry P, Mueller D, Osborne C, Elder A, Bomken SN, Pal D, Allan JM, Veal GJ, Cockerill PN, Wichmann C, Vormoor J, Lacaud G, Bonifer C, Heidenreich O. The Oncogenic Transcription Factor RUNX1/ETO Corrupts Cell Cycle Regulation to Drive Leukemic Transformation. Cancer Cell. 2018;34(4):626-42.e8.

57. Patrick R, Naval-Sanchez M, Deshpande N, Huang Y, Zhang J, Chen X, Yang Y, Tiwari K, Esmaeili M, Tran M, Mohamed AR, Wang B, Xia D, Ma J, Bayliss J, Wong K, Hun ML, Sun X, Cao B, Cottle DL, Catterall T, Barzilai-Tutsch H, Troskie RL, Chen Z, Wise AF, Saini S, Soe YM, Kumari S, Sweet MJ, Thomas HE, Smyth IM, Fletcher AL, Knoblich K, Watt MJ, Alhomrani M, Alsanie W, Quinn KM, Merson TD, Chidgey AP, Ricardo SD, Yu D, Jardé T, Cheetham SW, Marcelle C, Nilsson SK, Nguyen Q, White MD, Nefzger CM. The activity of early-life gene regulatory elements is hijacked in aging through pervasive AP-1-linked chromatin opening. Cell Metab. 2024;36(8):1858-81.e23.

58. Robinson MD, McCarthy DJ, Smyth GK. edgeR: a Bioconductor package for differential expression analysis of digital gene expression data. Bioinformatics. 2010 Jan 1;26(1):139–40.

59. Love MI, Huber W, Anders S. Moderated estimation of fold change and dispersion for RNA-seq data with DESeq2. Genome Biology. 2014;15(12):550.

60. Holmfeldt P, Ganuza M, Marathe H, He B, Hall T, Kang G, Moen J, Pardieck J, Saulsberry AC, Cico A, Gaut L, McGoldrick D, Finkelstein D, Tan K, McKinney-Freeman S. Functional screen identifies regulators of murine hematopoietic stem cell repopulation. J Exp Med. 2016;213(3):433–49.

61. Chen C, Yu W, Tober J, Gao P, He B, Lee K, Trieu T, Blobel GA, Speck NA, Tan K. Spatial Genome Reorganization between Fetal and Adult Hematopoietic Stem Cells. Cell Rep. 2019;29(12):4200-11.e7.

62. Unnikrishnan A, Papaemmanuil E, Beck D, Deshpande NP, Verma A, Kumari A, Woll PS, Richards LA, Knezevic K, Chandrakanthan V, Thoms JAI, Tursky ML, Huang Y, Ali Z, Olivier J, Galbraith S, Kulasekararaj AG, Tobiasson M, Karimi M, Pellagatti A, Wilson SR, Lindeman R, Young B, Ramakrishna R, Arthur C, Stark R, Crispin P, Curnow J, Warburton P, Roncolato F, Boultwood J, Lynch K, Jacobsen SEW, Mufti GJ, Hellstrom-Lindberg E, Wilkins MR, MacKenzie KL, Wong JWH, Campbell PJ, Pimanda JE. Integrative Genomics Identifies the Molecular Basis of Resistance to Azacitidine Therapy in Myelodysplastic Syndromes. Cell Reports. 2017;20(3):572–85.

63. Williams MS, Amaral FMR, Simeoni F, Somervaille TCP. A stress-responsive enhancer induces dynamic drug resistance in acute myeloid leukemia. The Journal of Clinical Investigation. 2020;130(3):1217–32.

64. Xu H, Muise ES, Javaid S, Chen L, Cristescu R, Mansueto MS, Follmer N, Cho J, Kerr K, Altura R, Machacek M, Nicholson B, Addona G, Kariv I, Chen H. Identification of predictive genetic signatures of Cytarabine responsiveness using a 3D acute myeloid leukaemia model. Journal of Cellular and Molecular Medicine. 2019;23(10):7063–77.

65. Zhang H, Nakauchi Y, Köhnke T, Stafford M, Bottomly D, Thomas R, Wilmot B, McWeeney SK, Majeti R, Tyner JW. Integrated analysis of patient samples identifies biomarkers for venetoclax efficacy and combination strategies in acute myeloid leukemia. Nature Cancer. 2020;1(8):826–39.

66. Aliee H, Theis FJ. AutoGeneS: Automatic gene selection using multi-objective optimization for RNA-seq deconvolution. Cell Syst. 2021;12(7):706-15.e4.

67. Eferl R, Wagner EF. AP-1: a double-edged sword in tumorigenesis. Nature Reviews Cancer. 2003;3(11):859–68.

68. Wang L, Gural A, Sun X-J, Zhao X, Perna F, Huang G, Hatlen MA, Vu L, Liu F, Xu H, Asai T, Xu H, Deblasio T, Menendez S, Voza F, Jiang Y, Cole PA, Zhang J, Melnick A, Roeder RG, Nimer SD. The Leukemogenicity of AML1-ETO Is Dependent on Site-Specific Lysine Acetylation. Science. 2011;333(6043):765–9.

69. Crowley SJ, Bednarski JJ, Magee JA, Li Y, White LS, Yang W. BCLAF1 Regulates Expression of AP-1 Genes and Fetal Hematopoietic Stem Cell Repopulation Activity. Blood. 2022;140(Supplement 1):2852–3.

70. Dell’Aversana C, Giorgio C, D’Amato L, Lania G, Matarese F, Saeed S, Di Costanzo A, Belsito Petrizzi V, Ingenito C, Martens JHA, Pallavicini I, Minucci S, Carissimo A, Stunnenberg HG, Altucci L. miR-194-5p/BCLAF1 deregulation in AML tumorigenesis. Leukemia. 2017;31(11):2315–25.

71. Tsherniak A, Vazquez F, Montgomery PG, Weir BA, Kryukov G, Cowley GS, Gill S, Harrington WF, Pantel S, Krill-Burger JM, Meyers RM, Ali L, Goodale A, Lee Y, Jiang G, Hsiao J, Gerath WFJ, Howell S, Merkel E, Ghandi M, Garraway LA, Root DE, Golub TR, Boehm JS, Hahn WC. Defining a Cancer Dependency Map. Cell. 2017;170(3):564-76.e16.

72. Asou H, Tashiro S, Hamamoto K, Otsuji A, Kita K, Kamada N. Establishment of a Human Acute Myeloid Leukemia Cell Line (Kasumi-1) With 8;21 Chromosome Translocation. Blood. 1991;77(9):2031–6.

73. Matozaki S, Nakagawa T, Kawaguchi R, Aozaki R, Tsutsumi M, Murayama T, Koizumi T, Nishimura R, Isobe T, Chihara K. Establishment of a myeloid leukaemic cell line (SKNO-1) from a patient with t(8;21) who acquired monosomy 17 during disease progression. Br J Haematol. 1995;89(4):805–11.

74. Körber V, Stainczyk SA, Kurilov R, Henrich KO, Hero B, Brors B, Westermann F, Höfer T. Neuroblastoma arises in early fetal development and its evolutionary duration predicts outcome. Nat Genet. 2023 Apr;55(4):619–630.

75. Cabezas-Wallscheid N, Eichwald V, de Graaf J, Löwer M, Lehr HA, Kreft A, Eshkind L, Hildebrandt A, Abassi Y, Heck R, Dehof AK, Ohngemach S, Sprengel R, Wörtge S, Schmitt S, Lotz J, Meyer C, Kindler T, Zhang DE, Kaina B, Castle JC, Trumpp A, Sahin U, Bockamp E. Instruction of haematopoietic lineage choices, evolution of transcriptional landscapes and cancer stem cell hierarchies derived from an AML1-ETO mouse model. EMBO Mol Med. 2013;5(12):1804–20.

76. Singh SK, Kagalwala MN, Parker-Thornburg J, Adams H, Majumder S. REST maintains self-renewal and pluripotency of embryonic stem cells. Nature. 2008;453(7192):223–7.

77. Dahl JA, Jung I, Aanes H, Greggains GD, Manaf A, Lerdrup M, Li G, Kuan S, Li B, Lee AY, Preissl S, Jermstad I, Haugen MH, Suganthan R, Bjørås M, Hansen K, Dalen KT, Fedorcsak P, Ren B, Klungland A. Broad histone H3K4me3 domains in mouse oocytes modulate maternal-to-zygotic transition. Nature. 2016;537(7621):548–52.

78. Nicosia L, Spencer GJ, Brooks N, Amaral FMR, Basma NJ, Chadwick JA, Revell B, Wingelhofer B, Maiques-Diaz A, Sinclair O, Camera F, Ciceri F, Wiseman DH, Pegg N, West W, Knurowski T, Frese K, Clegg K, Campbell VL, Cavet J, Copland M, Searle E, Somervaille TCP. Therapeutic targeting of EP300/CBP by bromodomain inhibition in hematologic malignancies. Cancer Cell. 2023;41(12):2136-53.e13.

79. Crowley S, White LS, Li Y, Yang W, Magee JA, Bednarski JJ. Bclaf1 promotes hematopoietic stem cell repopulating capacity and self-renewal. The Journal of Immunology. 2022;208(1_Supplement):47.01-47.01.

80. Wolf FA, Angerer P, Theis FJ. SCANPY: large-scale single-cell gene expression data analysis. Genome Biology. 2018;19(1):15.

81. Wolock SL, Lopez R, Klein AM. Scrublet: Computational Identification of Cell Doublets in Single-Cell Transcriptomic Data. Cell Systems. 2019;8(4):281-91.e9.

82. Heumos L, Schaar A, Consortium S-CBP. Single-cell best practices 2023. Available from: https://www.sc-best-practices.org/preamble.html#.

83. Büttner M, Miao Z, Wolf FA, Teichmann SA, Theis FJ. A test metric for assessing single-cell RNA-seq batch correction. Nature Methods. 2019;16(1):43–9.

84. Domínguez Conde C, Xu C, Jarvis LB, Rainbow DB, Wells SB, Gomes T, Howlett SK, Suchanek O, Polanski K, King HW, Mamanova L, Huang N, Szabo PA, Richardson L, Bolt L, Fasouli ES, Mahbubani KT, Prete M, Tuck L, Richoz N, Tuong ZK, Campos L, Mousa HS, Needham EJ, Pritchard S, Li T, Elmentaite R, Park J, Rahmani E, Chen D, Menon DK, Bayraktar OA, James LK, Meyer KB, Yosef N, Clatworthy MR, Sims PA, Farber DL, Saeb-Parsy K, Jones JL, Teichmann SA. Cross-tissue immune cell analysis reveals tissue-specific features in humans. Science. 2022;376(6594):eabl5197.

85. Aran D, Looney AP, Liu L, Wu E, Fong V, Hsu A, Chak S, Naikawadi RP, Wolters PJ, Abate AR, Butte AJ, Bhattacharya M. Reference-based analysis of lung single-cell sequencing reveals a transitional profibrotic macrophage. Nat Immunol. 2019;20(2):163–72.

86. Novershtern N, Subramanian A, Lawton LN, Mak RH, Haining WN, McConkey ME, Habib N, Yosef N, Chang CY, Shay T, Frampton GM, Drake AC, Leskov I, Nilsson B, Preffer F, Dombkowski D, Evans JW, Liefeld T, Smutko JS, Chen J, Friedman N, Young RA, Golub TR, Regev A, Ebert BL. Densely interconnected transcriptional circuits control cell states in human hematopoiesis. Cell. 2011 Jan 21;144(2):296–309.

87. Ianevski A, Giri AK, Aittokallio T. Fully-automated and ultra-fast cell-type identification using specific marker combinations from single-cell transcriptomic data. Nature Communications. 2022;13(1):1246.

88. Wu T, Hu E, Xu S, Chen M, Guo P, Dai Z, Feng T, Zhou L, Tang W, Zhan L, Fu X, Liu S, Bo X, Yu G. clusterProfiler 4.0: A universal enrichment tool for interpreting omics data. The Innovation. 2021;2(3).

