## Supplementary figures and tables for "Single-Cell Atlas of AML Reveals Age-Related Gene Regulatory Networks in t(8;21) AML"

**Supplementary Table 1 – Data Included in AML scAtlas**

| Study | DOI | Protocol | Samples Included | Data Source |
|---|---|---|---|---|
| Zheng et al, 2017 | https://doi.org/10.1038/ncomms14049 | 10X Genomics | 2 AML | http://support.10xgenomics.com/single-cell/datasets |
| Oetjen et al, 2018 | https://doi.org/10.1172/jci.insight.124928 | 10X Genomics | 25 Healthy BM (20 donors) | GSE120221 |
| Van Galen et al, 2019 | https://doi.org/10.1016/j.cell.2019.01.031 | Seq-Well | 16 AML<br>5 Healthy BM (4 donors) | GSE116256 |
| Setty et al, 2019 | https://doi.org/10.1038/s41587-019-0068-4 | 10X Genomics | 3 Healthy BM | https://explore.data.humancellatlas.org/projects/091cf39b-01bc-42e5-9437-f419a66c8a45 |
| Petti et al, 2019 | https://doi.org/10.1038/s41467-019-11591-1 | 10X Genomics | 5 AML | https://zenodo.org/records/3345981 |
| Pei et al, 2020 | https://doi.org/10.1158/2159-8290.cd-19-0710 | 10X Genomics (CITE-seq) | 1 AML | GSE143363 |
| Caron et al, 2020 | https://doi.org/10.1038/s41598-020-64929-x | 10X Genomics | 3 Healthy BM | GSE132509 |
| Jiang et al, 2020 | https://doi.org/10.1073/pnas.2003900117 | 10X Genomics | 9 AML | https://www.biosino.org/node/project/detail/OEP000629 |
| Johnston et al, 2020 | https://doi.org/10.1016/j.gene.2020.144758 | 10X Genomics | 2 AML (1 donor) | GSE145410 |
| Velten et al, 2021 | https://doi.org/10.1038/s41467-021-21650-1 | Muta-Seq | 4 AML | https://doi.org/10.6084/m9.figshare.12382685.v1 |
| Stetson et al, 2021 | https://doi.org/10.1038/s41375-021-01338-7 | Smart-Seq2 | 4 AML | GSE126068 |
| Zhai et al, 2022 | https://doi.org/10.1186/s12943-022-01635-4 | SORT-Seq | 6 AML | Provided by authors upon request |
| Human Cell Atlas, 2022 | NA | 10X Genomics | 16 Healthy BM (8 donors) | https://explore.data.humancellatlas.org/projects/cc95ff89-2e68-4a08-a234-480eca21ce79 |
| Lasry et al, 2023 | https://doi.org/10.1038/s43018-022-00480-0 | 10X Genomics | 42 AML<br>5 Healthy BM | GSE185381 |
| Li et al, 2023 | https://doi.org/10.1038/s41375-022-01789-6 | 10X Genomics | 4 AML | OMIX002180 |
| Naldini et al, 2023 | https://doi.org/10.1038/s41467-023-36969-0 | 10X Genomics | 16 AML | GSE185993 |
| Beneyto-Calabuig et al, 2023 | https://doi.org/10.1016/j.stem.2023.04.001 | 10X Genomics | 19 AML<br>1 Healthy BM | https://doi.org/10.6084/m9.figshare.20291628 |
| Zheng et al, 2023 | https://doi.org/10.1186/s13059-023-03031-7 | 10X Genomics | 13 AML<br>1 Healthy BM | Provided by authors upon request |
| Mumme et al, 2023 | https://doi.org/10.1038/s41467-023-41994-0 | 10X Genomics | 19 AML | GSE235923 |
| Fiskus et al, 2023 | https://doi.org/10.1038/s41408-023-00826-6 | 10X Genomics | 1 AML | GSE228326 |

### Supplementary Table 2 – Benchmarking Batch Correction Tools on AML scAtlas

| | Silhouette label | cLISI | Silhouette batch | iLISI | KBET | Graph connectivity | PCR comparison | Batch correction | Bio conservation | Total |
|---|---|---|---|---|---|---|---|---|---|---|
| scVI (Sample) - 2000 Genes | 0.906 | 0.7031 | 1 | 0.2446 | 0.6389 | 0.946 | 0.5011 | 0.8411 | 0.7899 | 1 |
| scVI (Sample) - 6000 Genes | 0.8582 | 0.7694 | 0.744 | 0.1903 | 0.5061 | 0.9801 | 0.5244 | 0.8202 | 0.798 | 0.9763 |
| scVI (Sample) - 10000 Genes | 0.8094 | 0.7579 | 0.7626 | 0.1976 | 0.5475 | 1 | 0.5095 | 0.8275 | 0.7644 | 0.9725 |
| scVI (Sample) - 4000 Genes | 0.8449 | 0.759 | 0.8628 | 0.1982 | 0.5065 | 0.96 | 0.5066 | 0.8042 | 0.7849 | 0.9507 |
| Harmony - 6000 Genes | 0.1891 | 0.1253 | 0.5579 | 0.9625 | 0.9112 | 0.1535 | 1 | 1 | 0.0795 | 0.9256 |
| Harmony - All Genes | 0.2296 | 0.0695 | 0.6557 | 0.9837 | 0.9638 | 0.1182 | 0.9933 | 0.9993 | 0.0729 | 0.9221 |
| Harmony - 4000 Genes | 0.2296 | 0.0695 | 0.6557 | 0.9837 | 0.9635 | 0.1182 | 0.9933 | 0.9992 | 0.0729 | 0.922 |
| Harmony - 10000 Genes | 0.1535 | 0.1487 | 0.5136 | 0.9459 | 0.8596 | 0.2086 | 0.987 | 0.9963 | 0.0717 | 0.9179 |
| Harmony - 8000 Genes | 0.1598 | 0.1468 | 0.5283 | 0.9538 | 0.868 | 0.1833 | 0.9912 | 0.9916 | 0.0743 | 0.9128 |
| scVI (Sample) - 8000 Genes | 0.7854 | 0.7704 | 0.7517 | 0.1899 | 0.2659 | 0.921 | 0.5165 | 0.7203 | 0.7574 | 0.8326 |
| Harmony - 2000 Genes | 0.2008 | 0 | 0.748 | 1 | 1 | 0 | 0.9698 | 0.9322 | 0.0198 | 0.8156 |
| scANVI (Sample) - 6000 Genes | 0.472 | 0.4303 | 0.4246 | 0.4866 | 0.7168 | 0.8284 | 0.5436 | 0.7872 | 0.4006 | 0.7788 |
| scVI (Sample) - All Genes | 0.663 | 0.7471 | 0.6518 | 0.2126 | 0.2492 | 0.9588 | 0.4679 | 0.673 | 0.6759 | 0.7403 |
| scANVI (Sample) - 8000 Genes | 0.4225 | 0.4515 | 0.5196 | 0.4611 | 0.6608 | 0.8141 | 0.5284 | 0.759 | 0.3838 | 0.7361 |
| scANVI (Sample) - 10000 Genes | 0.2636 | 0.45 | 0.4957 | 0.4731 | 0.7562 | 0.858 | 0.5059 | 0.7799 | 0.2933 | 0.7275 |
| scANVI (Sample) - All Genes | 0.3415 | 0.4354 | 0.4421 | 0.4828 | 0.7517 | 0.8291 | 0.5098 | 0.762 | 0.3295 | 0.7188 |
| scANVI (Sample) - 4000 Genes | 0.3495 | 0.3456 | 0.3818 | 0.5838 | 0.7254 | 0.7897 | 0.511 | 0.7318 | 0.2865 | 0.6633 |
| scVI (Study) - 2000 Genes | 0.9523 | 0.9128 | 0.6772 | 0.0335 | 0.4337 | 0.9299 | 0.2344 | 0.4533 | 0.927 | 0.5571 |
| scVI (Study) - 4000 Genes | 0.8783 | 0.9688 | 0.3721 | 0.0046 | 0.3356 | 0.9375 | 0.2394 | 0.4029 | 0.9148 | 0.4879 |
| scANVI (Sample) - 2000 Genes | 0 | 0.177 | 0 | 0.7712 | 0.955 | 0.7468 | 0.4155 | 0.628 | 0 | 0.4184 |
| scANVI (Study) - 4000 Genes | 0.5629 | 0.9246 | 0.1831 | 0.0313 | 0.3875 | 0.9418 | 0.2477 | 0.4068 | 0.7132 | 0.4142 |
| scVI (Study) - 10000 Genes | 0.8156 | 0.9745 | 0.2186 | 0.0037 | 0.202 | 0.9465 | 0.2307 | 0.3469 | 0.8823 | 0.4035 |
| scVI (Study) - 8000 Genes | 0.7385 | 0.9752 | 0.252 | 0.0035 | 0.1716 | 0.936 | 0.2441 | 0.3517 | 0.8392 | 0.3929 |
| scANVI (Study) - 8000 Genes | 0.609 | 0.9553 | 0.1597 | 0.0124 | 0.2222 | 0.957 | 0.2454 | 0.367 | 0.7555 | 0.3797 |
| scVI (Study) - 6000 Genes | 0.7372 | 0.9733 | 0.2688 | 0.004 | 0.062 | 0.9248 | 0.2508 | 0.3268 | 0.8374 | 0.3603 |
| scANVI (Study) - 6000 Genes | 0.5889 | 0.9429 | 0.1916 | 0.0176 | 0.2225 | 0.9434 | 0.2379 | 0.3555 | 0.7376 | 0.358 |
| scANVI (Study) - 10000 Genes | 0.6167 | 0.9512 | 0.1264 | 0.0135 | 0.1976 | 0.9027 | 0.2257 | 0.3075 | 0.7577 | 0.3044 |
| scANVI (Study) - All Genes | 0.6122 | 0.9612 | 0.0581 | 0.0086 | 0.1881 | 0.968 | 0.1719 | 0.2738 | 0.7604 | 0.2623 |
| scANVI (Study) - 2000 Genes | 0.3509 | 0.7896 | 0.1808 | 0.1386 | 0.5039 | 0.9054 | 0.1363 | 0.3006 | 0.5221 | 0.2034 |
| scVI (Study) - All Genes | 0.6792 | 0.9789 | 0.1341 | 0.0018 | 0 | 0.9609 | 0.144 | 0.2003 | 0.8076 | 0.1867 |
| Uncorrected - 2000 Genes | 0.7746 | 0.9893 | 0.7891 | 0.0256 | 0.498 | 0.7869 | 0 | 0.1559 | 0.867 | 0.1532 |
| Uncorrected - 4000 Genes | 0.9887 | 1 | 0.6323 | 0.002 | 0.2756 | 0.7349 | 0 | 0.0538 | 0.9936 | 0.0719 |
| Uncorrected - All Genes | 0.9887 | 1 | 0.6323 | 0.002 | 0.2714 | 0.7349 | 0 | 0.0527 | 0.9936 | 0.0706 |
| Uncorrected - 6000 Genes | 0.9919 | 1 | 0.5309 | 0.0002 | 0.3077 | 0.7116 | 0 | 0.0385 | 0.9954 | 0.0531 |
| Uncorrected - 8000 Genes | 1 | 1 | 0.5064 | 0 | 0.1888 | 0.7019 | 0 | 0.0003 | 1 | 0.0059 |
| Uncorrected - 10000 Genes | 0.9748 | 1 | 0.5019 | 0 | 0.1314 | 0.7304 | 0 | 0 | 0.9858 | 0 |

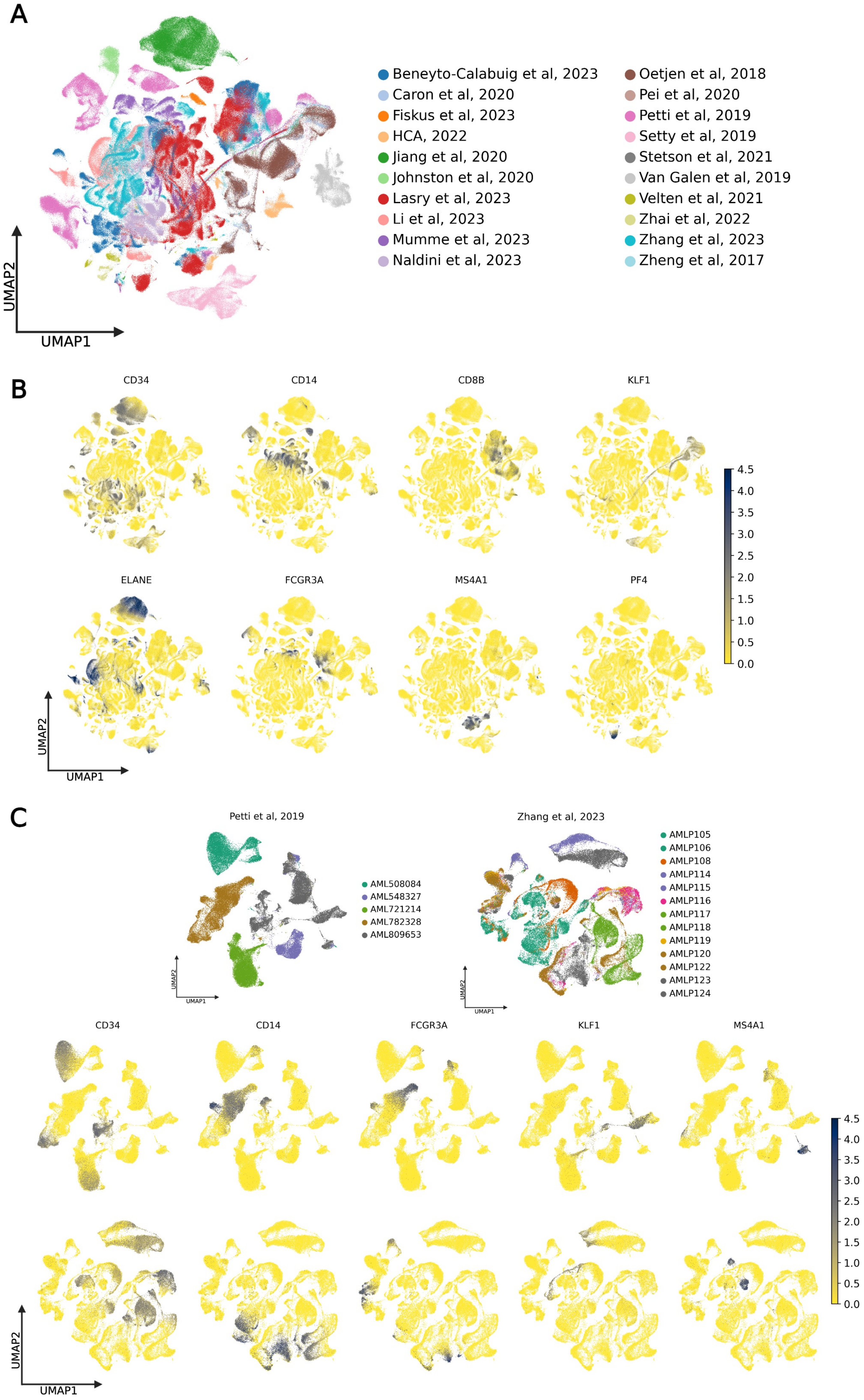
A
Beneyto-Calabuig et al, 2023
Caron et al, 2020
Fiskus et al, 2023
HCA, 2022
Jiang et al, 2020
Johnston et al, 2020
Lasry et al, 2023
Li et al, 2023
Mumme et al, 2023
Naldini et al, 2023
Oetjen et al, 2018
Pei et al, 2020
Petti et al, 2019
Setty et al, 2019
Stetson et al, 2021
Van Galen et al, 2019
Velten et al, 2021
Zhai et al, 2022
Zhang et al, 2023
Zheng et al, 2017
UMAP2
UMAP1
B
CD34
CD14
CD8B
KLF1
ELANE
FCGR3A
MS4A1
PF4
4.5
4.0
3.5
3.0
2.5
2.0
1.5
1.0
0.5
0.0
UMAP2
UMAP1
C
Petti et al, 2019
AML508084
AML548327
AML721214
AML782328
AML809653
Zhang et al, 2023
AMLP105
AMLP106
AMLP108
AMLP114
AMLP115
AMLP116
AMLP117
AMLP118
AMLP119
AMLP120
AMLP122
AMLP123
AMLP124
CD34
CD14
FCGR3A
KLF1
MS4A1
4.5
4.0
3.5
3.0
2.5
2.0
1.5
1.0
0.5
0.0
UMAP2
UMAP1

**Supplementary Figure 1 – Initial Analysis Establishes Presence of Batch Effects**

(**A**) Initial dimensionality reduction and UMAP plotting prior to batch correction of the 748,679 high quality cells. (**B**) Visualization of key hematopoietic marker genes on the uncorrected UMAP. (**C**) Representative study examples (Petti et al, 2019, and Zhang et al, 2023) investigating batch effects. UMAPs of different samples in each study (top panels), and hematopoietic marker genes (bottom panels; Petti et al top, Zhang et al bottom).

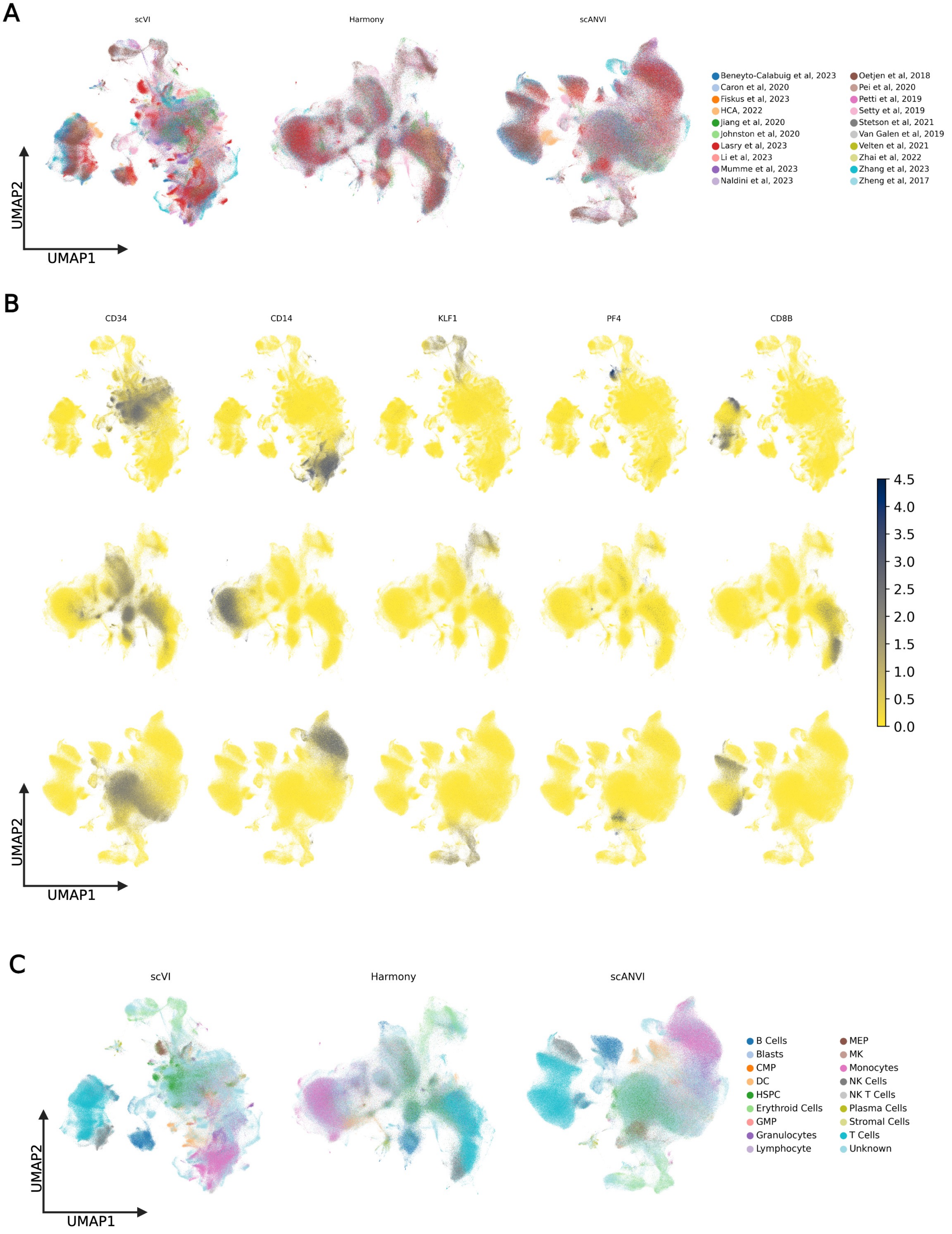
A
scVI
Harmony
scANVI
Beneyto-Calabuig et al, 2023
Caron et al, 2020
Fiskus et al, 2023
HCA, 2022
Jiang et al, 2020
Johnston et al, 2020
Lasry et al, 2023
Li et al, 2023
Mumme et al, 2023
Naldini et al, 2023
Oetjen et al, 2018
Pei et al, 2020
Petti et al, 2019
Setty et al, 2019
Stetson et al, 2021
Van Galen et al, 2019
Velten et al, 2021
Zhai et al, 2022
Zhang et al, 2023
Zheng et al, 2017
UMAP2
UMAP1
B
CD34
CD14
KLF1
PF4
CD8B
4.5
4.0
3.5
3.0
2.5
2.0
1.5
1.0
0.5
0.0
UMAP2
UMAP1
C
scVI
Harmony
scANVI
B Cells
Blasts
CMP
DC
HSPC
Erythroid Cells
GMP
Granulocytes
Lymphocyte
MEP
MK
Monocytes
NK Cells
NK T Cells
Plasma Cells
Stromal Cells
T Cells
Unknown
UMAP2
UMAP1

**Supplementary Figure 2 – Benchmarking Batch Correction Methods**

(**A**) Dimensionality reduction and UMAP visualization using the batch corrected embeddings for scVI (left), Harmony (middle) and scANVI (right) shows improved integration of different studies in all cases. (**B**) Visualization of hematopoietic marker genes on the UMAP for Harmony (top) and scANVI (bottom), shows improved harmonization of cell types following batch correction. (**C**) Projection of original publication cell type annotations, where available, onto UMAP plots for scVI (left), Harmony (middle) and scANVI (right).

**A**

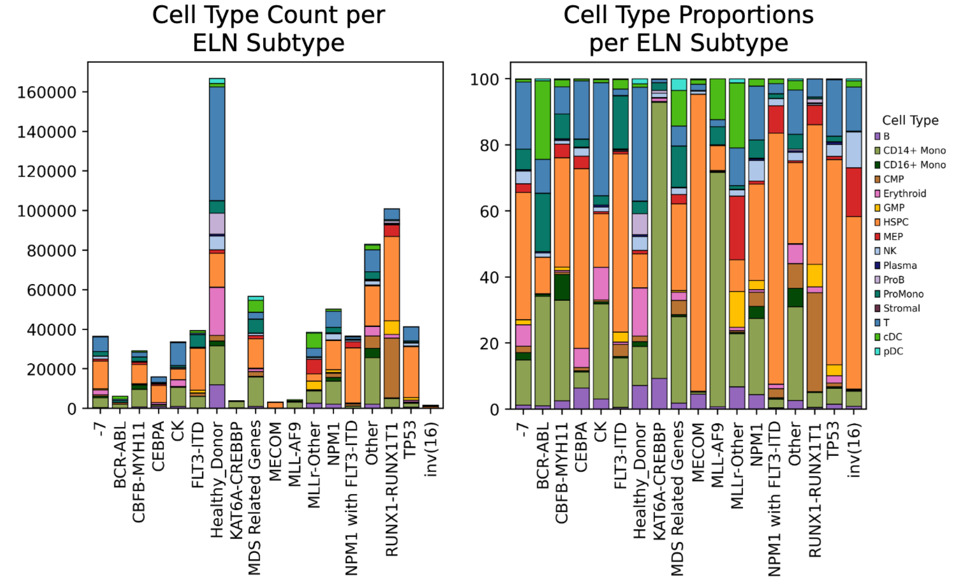


**Supplementary Figure 3 – Cell Type Proportions Vary by AML Subtype**

(**A**) Comparison of cell type abundance across different AML subtypes, shown as absolute values (left) and proportions (right).

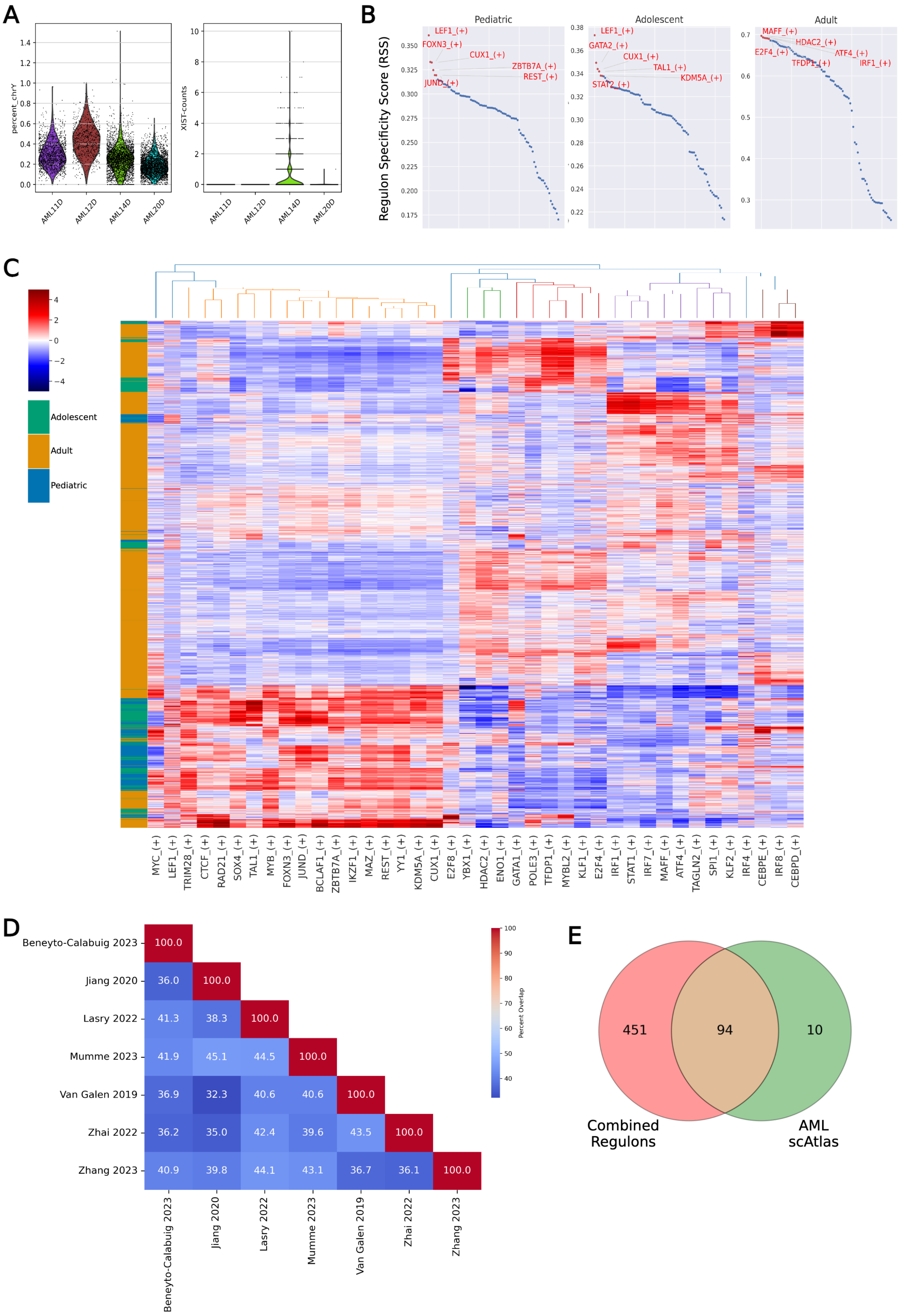
A
percent_chrY
XIST-counts
AML11D
AML12D
AML14D
AML20D
B
Regulon Specificity Score (RSS)
Pediatric
LEF1_(+)
FOXN3_(+)
CUX1_(+)
ZBTB7A_(+)
REST_(+)
JUND_(+)
Adolescent
LEF1_(+)
GATA2_(+)
CUX1_(+)
TAL1_(+)
KDM5A_(+)
STAT2_(+)
Adult
MAFF_(+)
HDAC2_(+)
E2F4_(+)
ATF4_(+)
TFDP1_(+)
IRF1_(+)
C
Adolescent
Adult
Pediatric
MYC_(+)
LEF1_(+)
TRIM28_(+)
CTCF_(+)
RAD21_(+)
SOX4_(+)
TAL1_(+)
MYB_(+)
FOXN3_(+)
JUND_(+)
BCLAF1_(+)
ZBTB7A_(+)
IKZF1_(+)
MAZ_(+)
REST_(+)
YY1_(+)
KDM5A_(+)
CUX1_(+)
E2F8_(+)
YBX1_(+)
HDAC2_(+)
ENO1_(+)
GATA1_(+)
POLE3_(+)
TFDP1_(+)
MYBL2_(+)
KLF1_(+)
E2F4_(+)
IRF1_(+)
STAT1_(+)
IRF7_(+)
MAFF_(+)
ATF4_(+)
TAGLN2_(+)
SPI1_(+)
KLF2_(+)
IRF4_(+)
CEBPE_(+)
IRF8_(+)
CEBPD_(+)
D
Beneyto-Calabuig 2023
Jiang 2020
Lasry 2022
Mumme 2023
Van Galen 2019
Zhai 2022
Zhang 2023
Percent Overlap
Beneyto-Calabuig 2023: 100.0
Jiang 2020: 36.0, 100.0
Lasry 2022: 41.3, 38.3, 100.0
Mumme 2023: 41.9, 45.1, 44.5, 100.0
Van Galen 2019: 36.9, 32.3, 40.6, 40.6, 100.0
Zhai 2022: 36.2, 35.0, 42.4, 39.6, 43.5, 100.0
Zhang 2023: 40.9, 39.8, 44.1, 43.1, 36.7, 36.1, 100.0
E
451
94
10
Combined Regulons
AML scAtlas

**Supplementary Figure 4 – AML with t(8;21) pySCENIC Analysis**

(**A**) XIST/ChrY expression in samples from patients with no recorded gender. (**B**) Regulon specificity scores (RSS) for pySCENIC regulons in each age group of the t(8;21) AML data analyzed. (**C**) Clustered heatmap of the AUC values (Z-score normalized) calculated using pySCENIC. After selecting for HSPCs, regulons were chosen based on their regulon specificity score (RSS). (**D**) Percentage overlap of regulon transcription factors (TFs) between individual studies with t(8;21) AML samples, when performing pySCENIC on each individually. (**E**) Overlap between the combined regulon TFs from individual study-wise iterations of pySCENIC and the integrated AML scAtlas dataset.

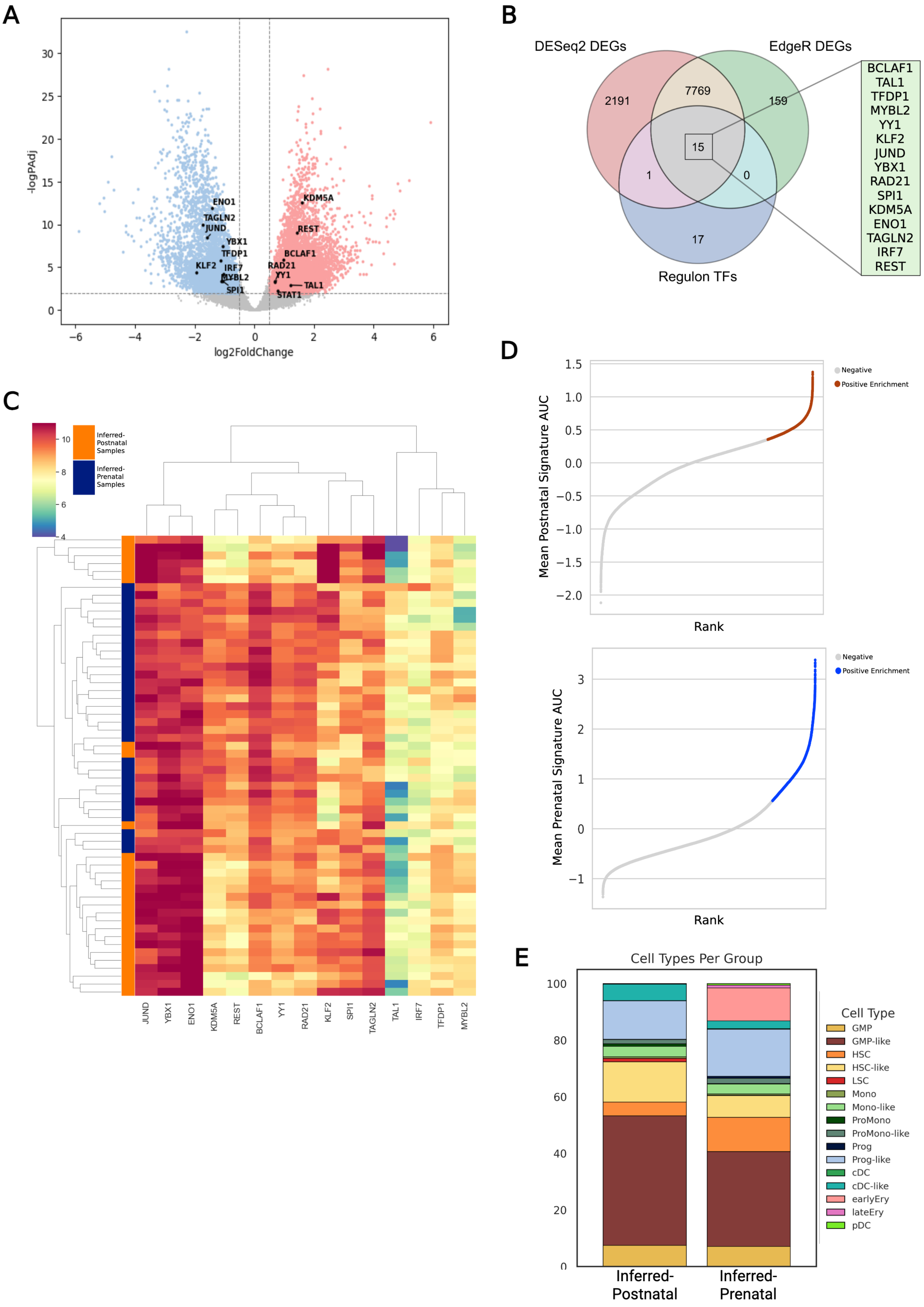
A
-logPAdj
log2FoldChange
ENO1
TAGLN2
JUND
YBX1
TFDP1
KLF2
IRF7
MYBL2
SPI1
KDM5A
REST
BCLAF1
RAD21
YY1
TAL1
STAT1
B
DESeq2 DEGs
EdgeR DEGs
Regulon TFs
2191
7769
159
15
1
0
17
BCLAF1
TAL1
TFDP1
MYBL2
YY1
KLF2
JUND
YBX1
RAD21
SPI1
KDM5A
ENO1
TAGLN2
IRF7
REST
C
Inferred-Postnatal Samples
Inferred-Prenatal Samples
JUND
YBX1
ENO1
KDM5A
REST
BCLAF1
YY1
RAD21
KLF2
SPI1
TAGLN2
TAL1
IRF7
TFDP1
MYBL2
D
Mean Postnatal Signature AUC
Rank
Negative
Positive Enrichment
Mean Prenatal Signature AUC
Rank
Negative
Positive Enrichment
E
Cell Types Per Group
Cell Type
GMP
GMP-like
HSC
HSC-like
LSC
Mono
Mono-like
ProMono
ProMono-like
Prog
Prog-like
cDC
cDC-like
earlyEry
lateEry
pDC
Inferred-Postnatal
Inferred-Prenatal

**Supplementary Figure 5 – Validation of pySCENIC Regulons**

(**A**) Differential gene expression volcano plot comparing the inferred-prenatal and inferred-postnatal bulk RNA-sequencing samples, as performed by DESeq2. (**B**) Venn diagram highlighting the intersect between regulon transcription factors (TFs), and two independent methods of differential gene expression. (**C**) Heatmap of regulon-associated TFs and their log normalized gene expression values across the samples in each group (prenatal origin versus postnatal origin). (**D**) Using the t(8;21) AML data from AML scAtlas, median absolute deviation (MAD) thresholding was used to select cells enriched for the inferred-prenatal origin (top) and inferred-postnatal origin (bottom) signatures. (**E**) Using the AML scAtlas cell type annotations from the HSPC/LSPC reference dataset, cell type proportions in the inferred-prenatal origin signature cells were compared to the postnatal origin cells.

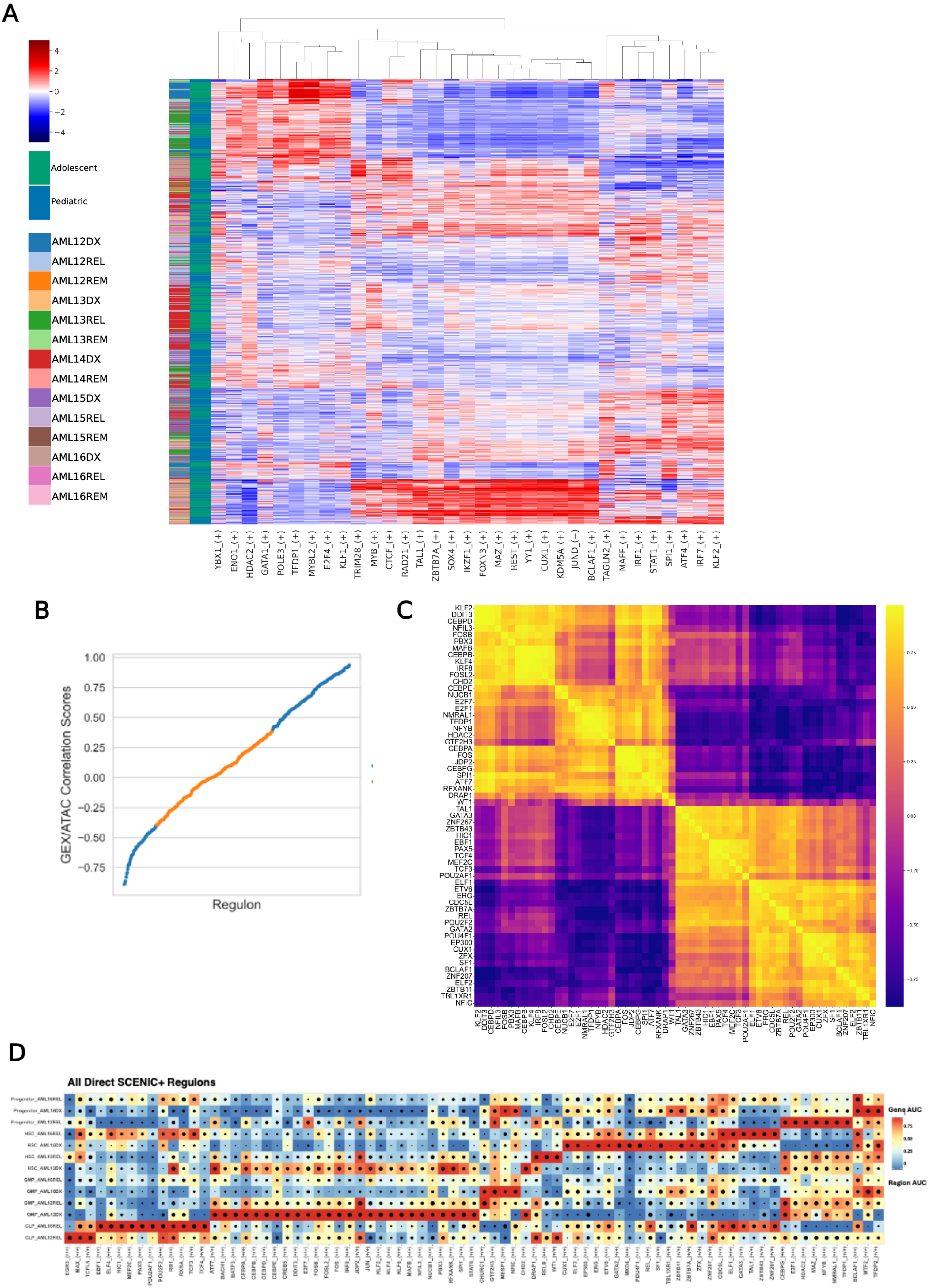
A
4
2
0
−2
−4
Adolescent
Pediatric
AML12DX
AML12REL
AML12REM
AML13DX
AML13REL
AML13REM
AML14DX
AML14REM
AML15DX
AML15REL
AML15REM
AML16DX
AML16REL
AML16REM
YBX1_(+)
ENO1_(+)
HDAC2_(+)
GATA1_(+)
POLE3_(+)
TFDP1_(+)
MYBL2_(+)
E2F4_(+)
KLF1_(+)
TRIM28_(+)
MYB_(+)
CTCF_(+)
RAD21_(+)
TAL1_(+)
ZBTB7A_(+)
SOX4_(+)
IKZF1_(+)
FOXN3_(+)
MAZ_(+)
REST_(+)
YY1_(+)
CUX1_(+)
KDM5A_(+)
JUND_(+)
BCLAF1_(+)
TAGLN2_(+)
MAFF_(+)
IRF1_(+)
STAT1_(+)
SPI1_(+)
ATF4_(+)
IRF7_(+)
KLF2_(+)
B
1.00
0.75
0.50
0.25
0.00
−0.25
−0.50
−0.75
GEX/ATAC Correlation Scores
Regulon
C
KLF2
DDIT3
CEBPD
NFIL3
FOSB
PBX3
MAFB
CEBPB
KLF4
IRF8
FOSL2
CHD2
CEBPE
NUCB1
E2F7
E2F1
NMRAL1
TFDP1
NFYB
HDAC2
GTF2H3
CEBPA
FOS
JDP2
CEBPG
SPI1
ATF7
RFXANK
DRAP1
WT1
TAL1
GATA3
ZNF267
ZBTB43
HIC1
EBF1
PAX5
TCF4
MEF2C
TCF3
POU2AF1
ELF1
ETV6
ERG
CDC5L
ZBTB7A
REL
POU2F2
GATA2
POU4F1
EP300
CUX1
ZFX
SF1
BCLAF1
ZNF207
ELF2
ZBTB11
TBL1XR1
NFIC
0.75
0.50
0.25
0.00
-0.25
-0.50
-0.75
D
All Direct SCENIC+ Regulons
Gene AUC
Region AUC

**Supplementary Figure 6 – Multiomics Data of t(8;21) AML Refines GRN**

(**A**) Using the Lambo et al dataset, age-associated regulon activity was calculated using pySCENIC AUCell. This identified patient samples highly enriched for the inferred-prenatal and inferred-postnatal origin signatures. (**B**) Plot showing the correlation scores between the scRNA-seq and scATAC-seq derived eRegulons. Correlation thresholds were used to prioritize eRegulons which correlate across modalities (highlighted in blue). (**C**) Correlation plot of eRegulon target genes shows clusters of related eRegulons with common targets. (**D**) SCENIC+ analysis identified a range of patient and cell type specific eRegulons. Dotplot shows all direct eRegulons inferred by SCENIC+ after initial filtering steps, split into patient associated cell type populations. This shows many eRegulons with a high correlation between scRNA-seq target gene activity (indicated by the color scale) and scATAC-seq target region accessibility (depicted by spot size).

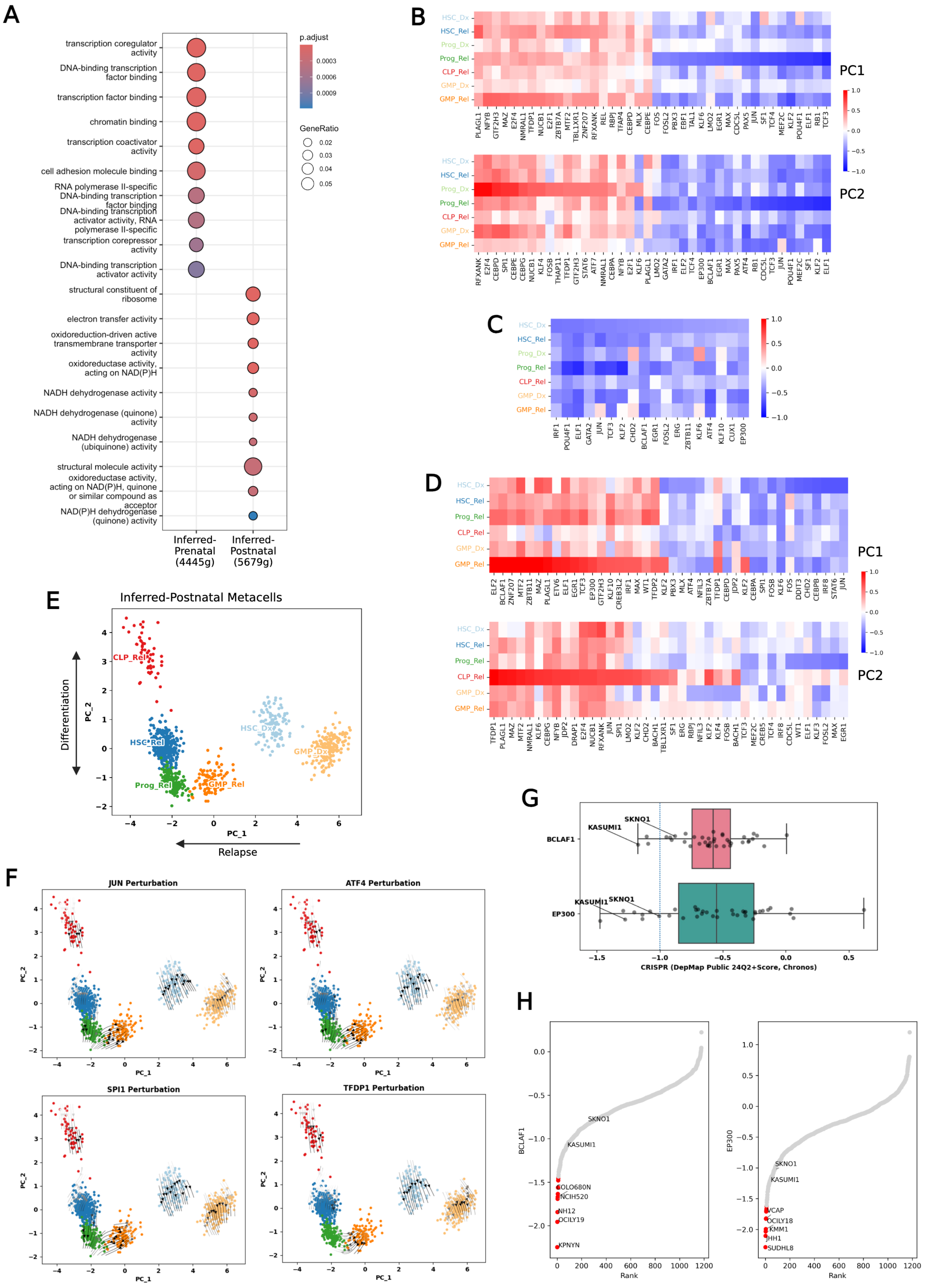
A
transcription coregulator activity
DNA-binding transcription factor binding
transcription factor binding
chromatin binding
transcription coactivator activity
cell adhesion molecule binding
RNA polymerase II-specific DNA-binding transcription factor binding
DNA-binding transcription activator activity, RNA polymerase II-specific
transcription corepressor activity
DNA-binding transcription activator activity
structural constituent of ribosome
electron transfer activity
oxidoreduction-driven active transmembrane transporter activity
oxidoreductase activity, acting on NAD(P)H
NADH dehydrogenase activity
NADH dehydrogenase (quinone) activity
NADH dehydrogenase (ubiquinone) activity
structural molecule activity
oxidoreductase activity, acting on NAD(P)H, quinone or similar compound as acceptor
NAD(P)H dehydrogenase (quinone) activity
Inferred-Prenatal (4445g)
Inferred-Postnatal (5679g)
p.adjust
0.0003
0.0006
0.0009
GeneRatio
0.02
0.03
0.04
0.05
B
HSC_Dx
HSC_Rel
Prog_Dx
Prog_Rel
CLP_Rel
GMP_Dx
GMP_Rel
PC1
PC2
C
D
PC1
PC2
E
Inferred-Postnatal Metacells
Differentiation
Relapse
CLP_Rel
HSC_Rel
Prog_Rel
GMP_Rel
HSC_Dx
GMP_Dx
PC_1
PC_2
F
JUN Perturbation
ATF4 Perturbation
SPI1 Perturbation
TFDP1 Perturbation
G
BCLAF1
EP300
KASUMI1
SKNO1
CRISPR (DepMap Public 24Q2+Score, Chronos)
H
BCLAF1
EP300
Rank
SKNO1
KASUMI1
OLO680N
NCIH520
NH12
OCILY19
KPNYN
VCAP
OCILY18
KMM1
JHH1
SUDHL8

**Supplementary Figure 7: SCENIC+ Analysis Identifies Age-Associated Candidate Perturbations in t(8;21) AML**

(**A**) Gene ontology over representation analysis of regulon target gene clusters, defined from the co-binding correlation map as inferred-prenatal or inferred-postnatal. GO molecular function gene sets used with an adjusted P value threshold of 0.05. (**B**) SCENIC+ perturbation simulation infers the predicted effect of knockout of selected TFs on the previously computed PCA embedding for the inferred-prenatal origin sample (AML16). Heatmap shows the predicted effect on PC1 (top) and PC2 (bottom) for the TFs with the largest predicted effect across all cell types. (**C**) SCENIC+ perturbation modelling results for the prenatal origin sample. Prioritized TFs based on predicted shift on the HSC compartment at diagnosis below -0.3. (**D**) Principal components analysis (PCA) of the gene based eRegulon enrichment scores for the inferred-postnatal origin samples at diagnosis and relapse. PC1 explains variance occurring between diagnosis and relapse. PC2 captures variance related to hematopoietic differentiation, split into myeloid and lymphoid trajectories. (**E**) SCENIC+ perturbation simulation results for the inferred-postnatal origin sample (AML12). Heatmap shows the predicted effect on PC1 (top) and PC2 (bottom) for the TFs with the largest predicted effect across all cell types. (**F**) SCENIC+ perturbation simulation shows the predicted effect of knockout of selected TFs on the previously computed PCA embedding. Arrows indicate the predicted shift in cell states relative to the initial PCA embedding. (**G**) DepMap CRISPR dependency scores for BCLAF1 (top) and EP300 (bottom) and as potential therapeutic targets identified in the prenatal t(8;21) AML sample, with relevant t(8;21) AML cell lines indicated. (**H**) DepMap CRISPR dependency scores for BCLAF1 (left) and EP300 (right), ranked for all cell lines.
